# Cryo-EM structures of human GPR155 elucidate its regulatory and transport mechanisms

**DOI:** 10.1101/2024.09.24.614577

**Authors:** Mansi Sharma, Dabbu K. Jaijyan, Sristi Nanda, Montserrat Samso, Wenhui Hu, Shikha Singh, Appu K. Singh

## Abstract

GPR155 is a polymodal lysosomal membrane “transceptor” comprising both a transporter domain and a GPCR domain, predominantly expressed in brain. GPR155 facilitates cholesterol-dependent mTORC1 signaling and is implicated in neurological disorders like Huntington’s disease. The GPCR domain likely does not bind extracellular ligands canonically, and the functional relationship between GPR155 domains remains unclear. Here, we report the first structures of monomeric human GPR155 and two distinct dimers, revealing an inward-open transporter domain and an inactivated GPCR domain with a unique luminal loop 7 conformation occupying orthosteric pocket. The dimeric assembly is cholesterol-sensitive: at low cholesterol, the transporter domain resembles plant PIN transporters and transports auxin molecules; at high cholesterol, it forms a unique dimer stabilized by cholesterol. Altogether, these findings have implications for uncovering new lysosomal signaling pathways.

## Introduction

Cholesterol and auxin (Indole-3-acetic acid, IAA) are two biomolecules that, while both essential, serve distinct functions across different types of organisms. Cholesterol is well-characterized in mammals and other lower organisms for its crucial functions in membrane integrity and fluidity, hormone synthesis, and lysosomal signaling (*1*). Conversely, auxin, a key plant hormone, is essential for plant growth and development, but its role in non-plant organisms is not well understood, primarily due to a lack of evidence related to auxin transporters, PIN-FORMED family proteins, in these systems (*2–6*). In humans, auxin is shown to be produced by gut microbiota as tryptophan metabolites and plays an important role in a number of physiological processes (*7, 8*). Its altered levels are implicated in various pathological conditions, including neurological disorders such as Huntington’s disease.

Within the cell, the lysosomes act as a central hub for cellular signaling, playing essential roles in cholesterol homeostasis and numerous signaling pathways (*9, 10*). Acting beyond its traditional role in macromolecule degradation, the lysosome surface serves as a signaling platform for cholesterol sensing and its metabolism (*11*). Cholesterol levels within the lysosome are tightly regulated, with fluctuations directly influencing various signaling pathways, including the activation of mTORC1 (*12*). Recently, the lysosomal membrane receptor GPR155, which is also known as the lysosomal cholesterol signaling (LYCHOS) protein, has been identified as crucial for cholesterol-dependent signaling in lysosomes. By binding cholesterol through its TM1 domain, GPR155 undergoes a change in its conformation that enables it to bind with the GTPase-activating protein complex GATOR1 (*12*). This interaction promotes Rag GTPase activation and subsequently leads to mTORC1 activation and localization at the lysosome.

GPR155 is shown to be widely expressed in several organs, including the stomach and brain. In the brain, it was shown to be highly expressed in the forebrain, particularly in the lateral striatum (*13*). In the lateral striatum of the mouse, the GPR155 expression profile was comparable to that of the CB1 (cannabinoid receptor type 1) and GAD1 (GABA-synthesizing enzyme, glutamate decarboxylase 1), suggesting similar roles in modulating sensorimotor and limbic inputs (*14*). Notably, the dysregulation and altered expression of GPR155 is associated with Huntington’s disease and autism spectrum disorders, paralleling the downregulation of CB1 receptors observed in the lateral striatum of transgenic Huntington’s disease mice (*14, 15*). The knockdown of a GPR155 homolog, Anchor, in *Drosophila melanogaster* resulted in thickened vein phenotypes and enlarged wing size, indicating its role in negatively regulating BMP signaling (*16*). GPR155 expression pattern correlates with several cancers and therefore it is an important biomarker: its downregulation is associated with hepatocellular carcinoma (*17*), gastric cancer (*18*), and thyroid carcinoma (*19*), while its upregulation is observed in UVR-induced mouse melanomas(*20*). Importantly, a mutation I357S in GPR155 has been linked to chemotherapy resistance in cancer patients(*21*).

The orphan receptor GPR155 architecture suggests that it represents a physiological concatenation that may function analogous to a transceptor by integrating features from two different protein families (*22, 23*). The distal C-terminus of GPR155 contains a DEP domain, common in signaling proteins such as Regulator of G-protein Signaling (RGS) proteins, which increase the GTPase activity of G-protein (*24, 25*). This DEP domain in RGS7 proteins is known to interact with a class C family receptor GPR158, partially overlapping with G-protein binding sites (*26, 27*). However, the functional implications of DEP domain in GPR155 are not well understood. Moreover, the molecular architecture and functional organization of this lysosomal transceptor remain elusive. Here, we provide the first structural insights into the physiological concatemer GPR155, where the transporter domain exhibits structural similarities with plant auxin transporter proteins from the PIN family and assembles with a class B GPCR domain. These structures reveal new insights into the potential roles of auxin and cholesterol signaling which have implications for lysosome regulation in physiology and pathophysiology.

### Structure determination of GPR155

Human GPR155 is evolutionarily conserved from insects to humans (fig. S1) and comprises a transporter domain fused with a GPCR domain. When GPR155 is heterologously expressed in HEK293T cells, it specifically localizes to the lysosomes with no detectable expression in the plasma membrane or other organelles, corroborating previous studies (*12*) (fig. S2A). To elucidate the molecular details of functional coupling between the transporter and GPCR domains, we heterologously expressed GPR155 in suspension-adapted HEK293F cells using a baculovirus-based expression system and conducted Fluorescence Size Exclusion Chromatography (FSEC) screening in various detergents supplemented with lipids to optimize its biochemical behavior for structural studies. Interestingly, FSEC analysis indicated that cholesteryl hemisuccinate (CHS), a synthetic analog of cholesterol, plays a crucial role in determining the oligomeric state of detergent-solubilized GPR155. In the absence of CHS, we observed two distinct FSEC peaks: one slightly larger, leftward peak representing the dimeric assembly and another relatively smaller peak shifted to the right, indicating the monomeric state. However, in the presence of CHS, a predominant FSEC peak corresponding to a monomeric assembly, with minimal evidence of a dimeric form and even size exclusion purification of GPR155 indicated a similar trend (fig. S2C and D). Notably, the inclusion of CHS during GPR155 solubilization significantly increased the thermostability of protein relative to LMNG detergent alone (fig. S2B). Previously, GPR155 has been shown to function as a cholesterol sensor (*12*), our finding indicates that the mechanism by which GPR155 senses cholesterol could be related to its oligomeric state. As GPR155 exhibited relatively more stability in the presence of CHS, we purified GPR155 to homogeneity using lauryl maltose neopentyl glycol (LMNG) detergent supplemented with CHS for structural characterization using cryo-electron microscopy (cryo-EM). Initial reference-free 2D class averages revealed substantial structural heterogeneity and different types of assemblies (fig. S3B). Following three-dimensional classification and subsequent homogeneous and non-uniform refinements, we obtained cryo-EM structures of the GPR155 monomer at a resolution of 3.9 Å and the two dimeric forms at resolutions of 3.48 Å and 3.8 Å, respectively (Fig. 1, figs. S3, S4 and S5). The maps enabled de novo building of the entire protein structure using an AlphaFold model as a reference, with the exception of residues 1-32 from the luminal N-terminus and a long intracellular loop from residues 548-650, likely due to higher flexibility in these regions.

**Figure 1.**
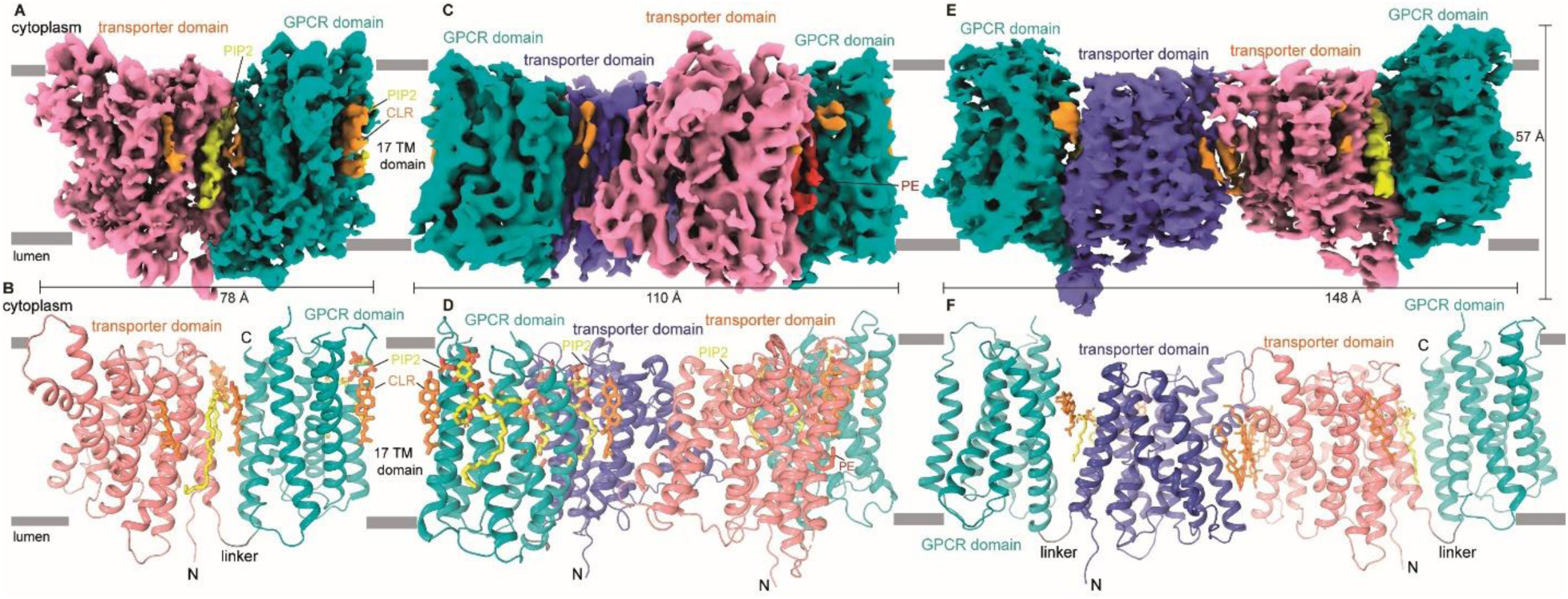
Overall architecture of GPR155 transceptor. **(A)** cryo-EM map the monomeric form of GPR155, with the transporter domain and GPCR domain shown in salmon and teal colors, respectively. **(B)** Cartoon representation of the side view of GPR155 in its monomeric form colored as in A. **(C)** cryo-EM map of the PIN-like dimeric arrangement of GPR155, with the transporter domains from the two monomeric subunits shown in purple and salmon colors, forming the dimeric interface, and GPCR domains shown in teal color. **(D)** Cartoon representation of the side view of GPR155 in its PIN-like dimeric form colored as in the C panel. Cholesterol (orange) and lipid moieties PIP2 (yellow) and PE (red) lined the cavity between the transporter and GPCR domains in the monomeric form. **(E)** cryo-EM map of the elongated cholesterol-driven dimer of GPR155. **(F)** Cartoon representation of the side view of GPR155 in its cholesterol-driven dimeric form colored as in E.

Interestingly, GPR155 dimer formation occurs exclusively via an inter-transporter domain interface. A closer examination revealed that different regions of the transporter domain participate in forming distinct dimeric interfaces. One dimer exhibits an arrangement reminiscent of plant auxin transporter dimers that belong to the PIN protein family (*3–5*), we have termed this “PIN-like dimer” (Fig. 1C and D, fig. S6, D, E, and F). In contrast, the other dimer has a more elongated arrangement of its two protomers, primarily stabilized by six cholesterol molecules embedded at the dimeric interface, which we refer to as the “cholesterol-driven dimer” (Fig. 1 E and F, fig. S6 G, H, and I).

### GPR155 transceptor architecture and domain organization

The monomeric structure of GPR155 offers insights into physiological transmembrane protein concatenation, where a transporter domain is fused to a GPCR domain, resulting in a unique architecture. The transmembrane domain (TMD) of GPR155 consists of 17 transmembrane helices (17 TM) connecting intracellular (ICL1-8) and luminal loops (LL1-8) (*12*). The N-terminus protrudes into the lumen of the lysosome, while its C-terminus extends into the cytoplasm (Fig. 1 and 2). Transmembrane helices 1-10 fold into a transporter-like structure that adopts a NhaA fold, similar to the PIN family that includes the auxin efflux transporters in plants (*3–5*), bacterial Na^+^/H^+^ antiporters (*28*), Na^+^/H^+^ exchangers (*29*) such as SLC9A, human solute carriers such as SLC10A, and apical sodium-dependent bile acid transporter (ASBT) homologs like ASBT_NM_ and ASBT_YF_ (Fig. 2, fig. S7A-D, H) (*30, 31*). The C-terminal TM helices 11-17, adopt a fold reminiscent of the GPCR family of proteins and exhibit close similarity to the Class B GPCRs (Fig. 2A, B, E-G, and fig. S8).

**Figure 2.**
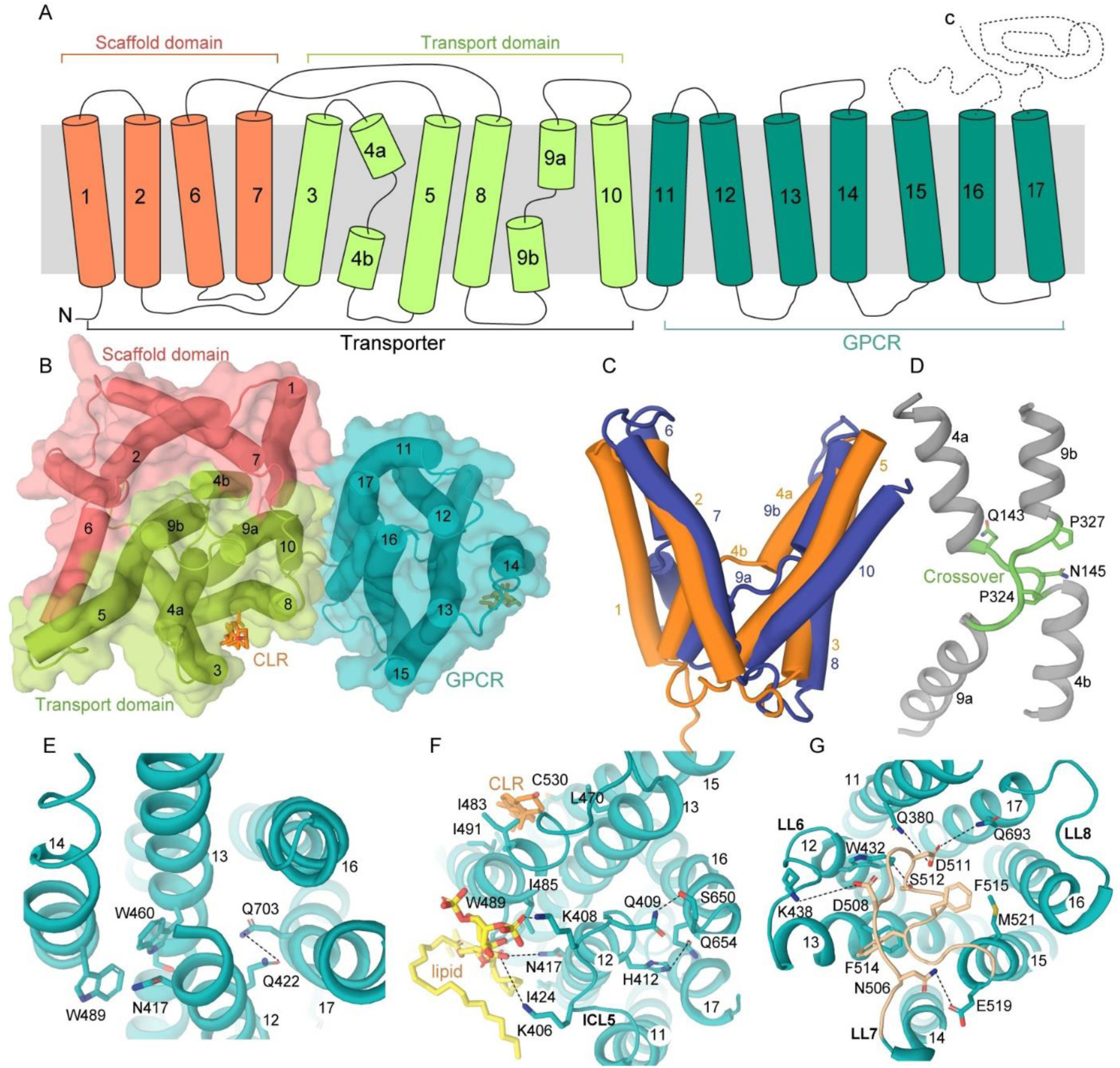
Domain architecture. **(A)** Domain organization diagram of the GPR155 subunit, comprising the transporter domain and GPCR domain (teal). The two subdomains found transporter domain: the scaffold domain (orange) at the N-terminus, followed by the transport domain (light green). Helices are shown as cylinders numbered 1-17, with interconnecting loops depicted as solid black lines, and the unresolved loop regions shown as dashed lines. **(B)** Cytoplasmic view of GPR155 with labeled domains. **(C)** Superimposed inverted repeats of TM1-TM5 and TM6-TM10. **(D)** A view of the central crossover (green) between TM4 and TM9. **(E)** Class B GPCR conserved motif residues shown in stick from TM14 and TM17, establishing interactions to stabilize the 7TM bundles. **(F)** Cytoplasmic interface of 7TM bundle. **(G)** Luminal interface of the 7TM bundle with the LL7 loop occupying the ligand-binding orthosteric site, shown in wheat color.

Within the transporter domain, TM1-5 and TM6-10 exhibit a pseudo-two-fold symmetry, forming two superimposable inverted repeats with a root mean square deviation (rmsd) of 5.4 Å for 160 Cα atoms (Fig. 2C) (*32*). The overall structure of the transporter domain can be split into two subdomains: a scaffold subdomain (TM1-2 and TM6-7) and a transport subdomain (TM3-5 and TM8-10) (Fig. 2A, B). Within the transport subdomain, TM4 and TM9 are discontinuous, forming two half-helices joined by the highly conserved sequences 143-QSND-146 and 321-GVFPVAP-327, respectively (Fig. 2A, D, fig. S1). These unwound helices cross over in an “X” shape at the center of the transmembrane domain (TMD), a structural feature also observed in the NhaA fold (*29*), in members of the PIN family (*3–5*), and Na^+^/H^+^ exchangers (Fig. 2D) (*28*). This signature helical crossover is shown to be broadly present in many transporters and plays a crucial role in substrate coordination and transport (*33–36*). In GPR155, the X-shaped crossover from TM4 contains a highly conserved N145 residue (fig. S1), an analogous asparagine residue is shown to directly bind with the indole-3-acetic acid (IAA) and *N*-1-naphthylphthalamic acid (NPA) in the PIN family of auxin transporters in plants (fig. S7f and G) (*3–5*), suggesting its involvement in substrate binding. Within, the X-shaped crossover, another conserved residue is contributed from TM9, P324, and mutating the analogous proline in the AEC transporter with other amino acids significantly impaired transporter function, further implicating these conserved residues in transceptor-mediated trafficking and substrate transport (*37*).

The C-terminal transmembrane helices (TM11-TM17) of GPR155 fold into a 7TM domain structure that closely resembles the typical architecture of G protein-coupled receptors (GPCRs)(*38, 39*). At the amino acid level, GPR155 exhibited very low sequence identity (18-20%) with class A and class B GPCRs. However, structure-based sequence alignment revealed a highly conserved “488-GWGxP-492” motif in TM14 (analogous to TM4 in GPCRs), allowing us to classify this GPCR domain as belonging to the class B family (figs. S1 and S8A) (*40*). Additionally, we found another class B family motif, “QG,” in TM17, further supporting this classification (figs. S1 and S8A). These two conserved motifs play important structural roles in stabilizing the overall configuration of TM12, TM13, TM14, and TM17 through key interactions. The G488^4.49b/4.49^ in the first conserved motif induces a bulge in TM14, resulting in crucial positioning of W489^4.50b/4.50^ in close proximity to TM12 and TM13, where it forms a hydrogen bond with the side chain of N417 from TM12 and also appear to engage in π-π stacking with W460 from TM13, similar to class B receptors such as the glucagon receptor (*41*) (Fig. 2E). Furthermore, the Q703 in the second motif present in TM17 interacts with Q422 from TM12, further providing overall stability to the 7TM bundle (Fig. 2E).

This C-terminal 7TM bundle is connected to the N-terminal transporter domain via a short three-amino-acid linker (residues 369-TMD-371) (Fig. 1B). The overall architecture of the 7TM helical bundle comprises three intracellular loops (ICL6-ICL8) and three luminal loops (LL6-LL8), which are analogous to the ICL1-ICL3 and ECL1-ECL3 loops found in classical GPCRs. The stability of the 7TM bundle is maintained by a series of hydrophobic and hydrophilic interactions at both the intracellular and luminal interfaces. On the intracellular side, the 7TM interface is characterized by both hydrophilic and hydrophobic interactions involving intracellular loop 6 (ICL6) and transmembrane segments TM12, TM13, TM16, and TM17 (Fig. 2F). The first layer of interactions includes a pair of hydrogen bonds between Q409 in ICL5 and S650 in TM16, while H412 in TM12 forms a hydrogen bond with Q654 in TM16, contributing to the intracellular interface of the 7TM bundle. Additionally, a cluster of basic residues—K405, K406, and K408— in ICL5 binds to a negatively charged head group of lipid-like molecules with its hydrophobic tail loosely packed against hydrophobic residues from TM13 and TM14, further enhancing the structural stability of the protein. Additionally, a cholesterol molecule is observed to be well-ordered within a hydrophobic cavity formed by residues in TM13 (L459, I466, L470), TM14 (I483, I491), and TM15 (C530, I534), potentially conferring additional stability to the receptor (Fig. 2F).

The luminal interface of the 7TM bundle is formed by interactions between transmembrane segments TM11-13, TM15, TM17, and the luminal loops LL6 and LL7 (Fig 2G and fig. S8), and is discussed in the next section. The C-terminal region of the GPCR is unique in that it includes an unusually large intracellular loop 8 (ICL8) between TM15 and TM16, which is analogous to ICL3 in typical GPCRs. However, in the GPR155 structure, this region is disordered, likely due to the inherent flexibility associated with these loop regions. The distal C-terminus folds into a Disheveled, Egl-10, and Pleckstrin (DEP) domain, which is similarly disordered, reflecting the dynamic nature of these regions within the overall protein structure.

### GPR155 luminal loop7 (LL7) occupies orthosteric ligand binding pocket

The structure of GPR155 features an 18-aminoacid-long luminal loop (LL7) within the GPCR domain, analogous to ECL2 in classical GPCRs, adopting a unique conformation reminiscent of constitutively active class A GPCRs like GPR52, GPR21, GPR101, and GPR161 (Fig. 2G, fig. S8B, D-F) (*42–45*). Structural superimposition of the GPR155 GPCR domain with GPR52 reveals that LL7 occupies a position similar to the self-activating ECL2 and occludes the canonical orthosteric ligand-binding pocket (fig. S8F). Unlike class A GPCRs, this arrangement of LL7 within the large hydrophobic cavity stabilizes the GPCR domain in its inactivated conformation as evidenced by structural superimposition with inactivated glucagon receptor (fig. S8D). This structural superimposition revealed that TM16 of GPR155 did not exhibit outward movement, a structural feature of inactivated GPCRs (fig. S8D). Furthermore, comparison with the ligand-bound state of class B GPCRs such as glucagon receptors indicated that LL7 occludes the ligand-binding pocket, with the residues that typically stabilize the helical peptide instead stabilizing the LL7 loop (*41*) (fig. S8E). This suggests an agonist-like, self-activating conformation of LL7, first discovered in the GPR52 receptor of the Class A family (*42*). At the apex of these interactions, a salt bridge forms between D508 in LL7 and K438 in LL6, while a hydrogen bond between N506 and E519 serves as a key stabilizing interaction. Additionally, the side chain of D511 in LL7 interacts with Q380 in TM11, and another hydrogen bond forms between S512 in LL7 and W432 in TM12, further strengthening the interface (Fig. 2G). The LL7 activating conformation is further stabilized by a hydrogen bond between D511 in LL7 and Q693 in TM17. Alongside these hydrophilic interactions, hydrophobic contacts also contribute to the stability of the LL7 region. Specifically, residues F515 and F514 in LL7 interact with F447 in TM13 and M521 in TM15, respectively. These combined hydrophilic and hydrophobic interactions create a structurally robust luminal interface, with LL7 acting as a lid that effectively seals the luminal, inverted V-shaped ligand-binding cavity (Fig. 2E).

### Role of lipids in the transporter and GPCR concatenation

The assembly of the concatemer is driven by helices TM1, TM7, TM8, and TM10 from the transporter, which pack against TM11, TM16, and TM17 from the GPCR (Fig. 3). These helices do not make extensive amino acid contacts but create a positively charged environment for the binding of phospholipids. The primary driving force for the concatemer assembly is a pair of deeply buried phospholipids, PIP2 and PE, at the interface of the transporter and GPCR.

**Figure 3.**
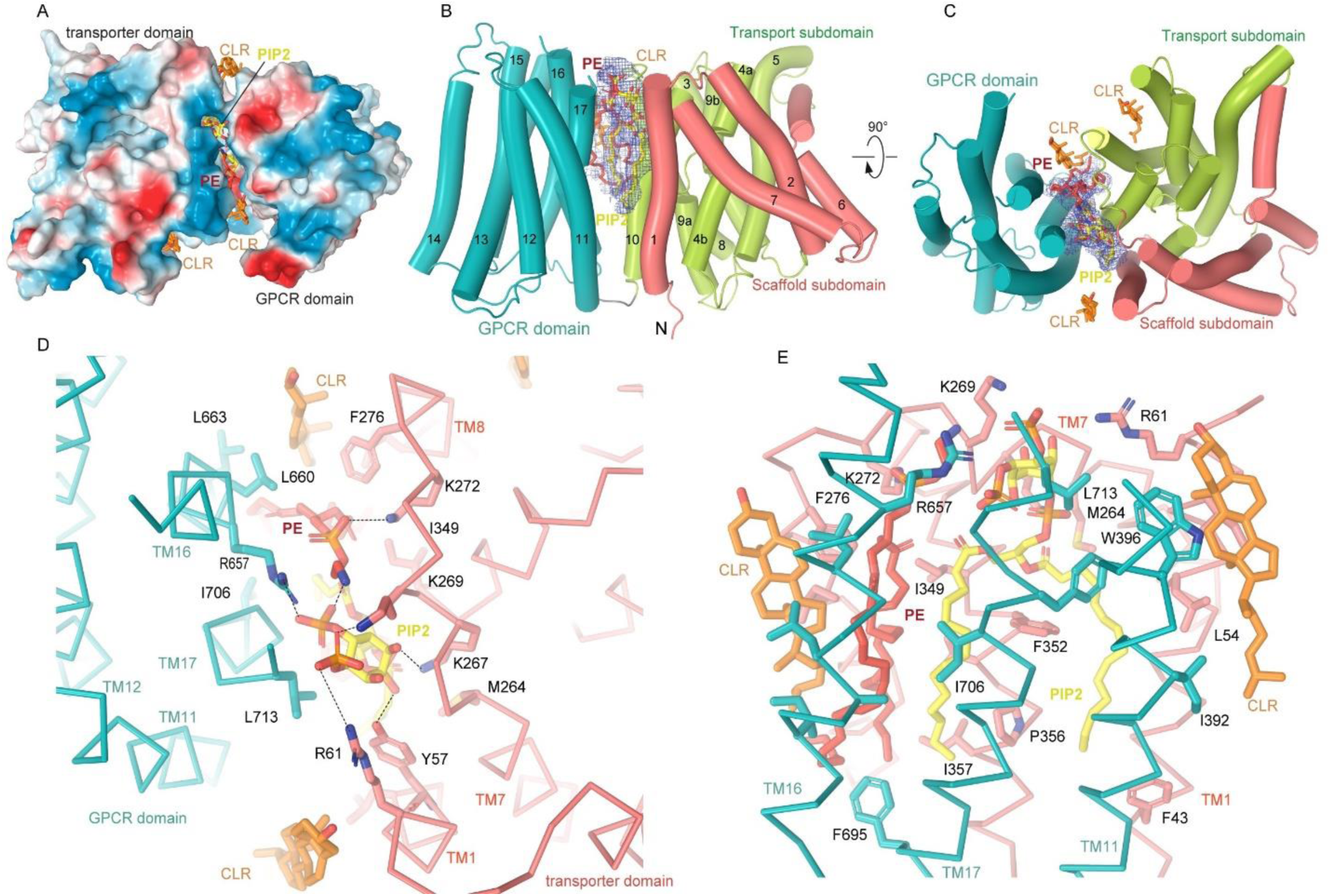
Lipids stabilize the concatemer assembly. **(A)** Electrostatic surface representation of the GPR155 protein. Positive charges are shown in blue, negative charges in red, and hydrophobic regions in white. The lipid molecules buried at the domain interface include PIP2 (yellow), PE (red), and cholesterol (orange), which are depicted in stick representation. **(B and C)** Two orthogonal views of the GPR155 monomer, with helices shown as cylinders. Cholesterol and lipid moieties are displayed as sticks, with their densities shown as blue mesh. **(D and E)** The hydrophobic and hydrophilic residues interacting with PIP2, PE, and cholesterol are shown as sticks.

The negatively charged head groups of these lipids are nestled in the positively charged cytoplasmic interface between the transporter and GPCR (Fig. 3 A-C). The negatively charged phosphate groups form extensive interactions with positively charged amino acids and polar contacts. Specifically, phosphate 1 of PIP2 is stabilized by Y57, R61, and K267 from the transporter, while phosphate 2 interacts with K269, and phosphate 3 is stabilized by R270 from the transporter and R657 from the GPCR. Similarly, PE is stabilized by K272 from the transporter and R270 and Q653 from the GPCR (Fig. 3C). These positively charged amino acids are fairly conserved from insects to human, suggesting a similar role in other GPR155 homologs (fig. S1). To understand the role of these positively charged stretches of amino acids in transceptor assembly, we substituted K269A, K273A, in addition to two double mutants K269A-R270A and K272A-K273A. These mutations did not affect trafficking of transceptor or the oligomeric assembly of GPR155, suggesting that other amino acids that interact with phospholipids, including hydrophilic and hydrophobic residues are also important for stabilizing the lipids at the domain interface (fig. S9). The hydrophobic tails of PIP2 and PE are sandwiched between the GPCR and transporter, forming extensive hydrophobic interactions with the hydrophobic amino acids (Fig. 3D). Specifically, the acyl chain of PIP2 forms hydrophobic contacts with residues from the GPCR domain, including V392 and W396 from TM11, L663 and I367 from TM16, and I706 and F708 from TM17. In the transporter domain, the acyl chain of PIP2 interacts with F43 from TM1, as well as I349, F352, V352, and P356 from TM10. Similarly, the acyl chain of PE stacks against hydrophobic residues I667 from TM16, F695 from TM17, and F276 and I280 from TM8 (Fig. 3E). Additionally, a pair of cholesterol molecules—one located between TM1 and TM17, and the other between TM8 and TM11 at the GPCR-transporter interface—also contribute towards stabilizing the concatemer assembly (Fig. 2F and Fig. 3E). Notably, the cholesterol bound to TM1 and TM17 is proposed to play a role in cholesterol sensing and its impact on GATOR1 interactions (*12*).

### Dimeric assembly and subunit interfaces

GPR155 was proposed as a cholesterol sensor, which plays an important role in GATOR1-dependent mTORC1 signaling in lysosomes (*12*). Specifically, cholesterol was shown to interact with TM1, resulting in enhanced coupling of GPR155 with the GATOR1 complex, suggesting that cholesterol binding can alter its physiological function. We obtained structures of two distinct dimers for GPR155: PIN-like dimer and cholesterol-driven dimer (Fig. 1, fig. S6 D-F), both of which are formed exclusively by the transporter region of monomers while the GPCR domain is distal to the dimeric interface. In the PIN-like dimer, dimerization is primarily driven by two-fold symmetric interactions between transmembrane helices TM1–2, TM7, and the linker connecting TM1 and TM2 within the scaffold subdomains of the two transporter domains, similar to that observed in plant auxin transporters (*3–5*) (Fig. 4A, fig. S7). These symmetric interactions involve extensive hydrophobic and hydrophilic contacts with precise helix alignment, creating a stable dimer interface. TM1 and TM2 from one subunit interact with TM7 and the corresponding helices from the other subunit, forming a symmetrical arrangement that maximizes the interaction surface area and stabilizes the overall dimeric structure. At the cytoplasmic interface, a pair of symmetric hydrogen bonds is formed between the side chains of N63 and R79 from both protomers. Additionally, a crucial hydrogen bond is established between the backbone carbonyl of I65 from the TM1–TM2 linker of subunit A and the side chain of N75 from TM2 of subunit B (Fig. 4B). Additionally, the side chain of F258 from subunit A stacks against F76 and F80 from TM1 of subunit B (Fig. 4C and D). Further, E48 from the luminal half of TM1 engages in another hydrogen bond with N243, strengthening this dimeric interface (Fig. 4D). These interactions ensure the proper orientation and proximity of the helices, contributing to the stability of the dimer. In addition to hydrophilic contacts, the hydrophobic symmetric interaction between F258 and L248 from TM7 of both protomers further strengthens the dimeric interface.

**Figure 4.**
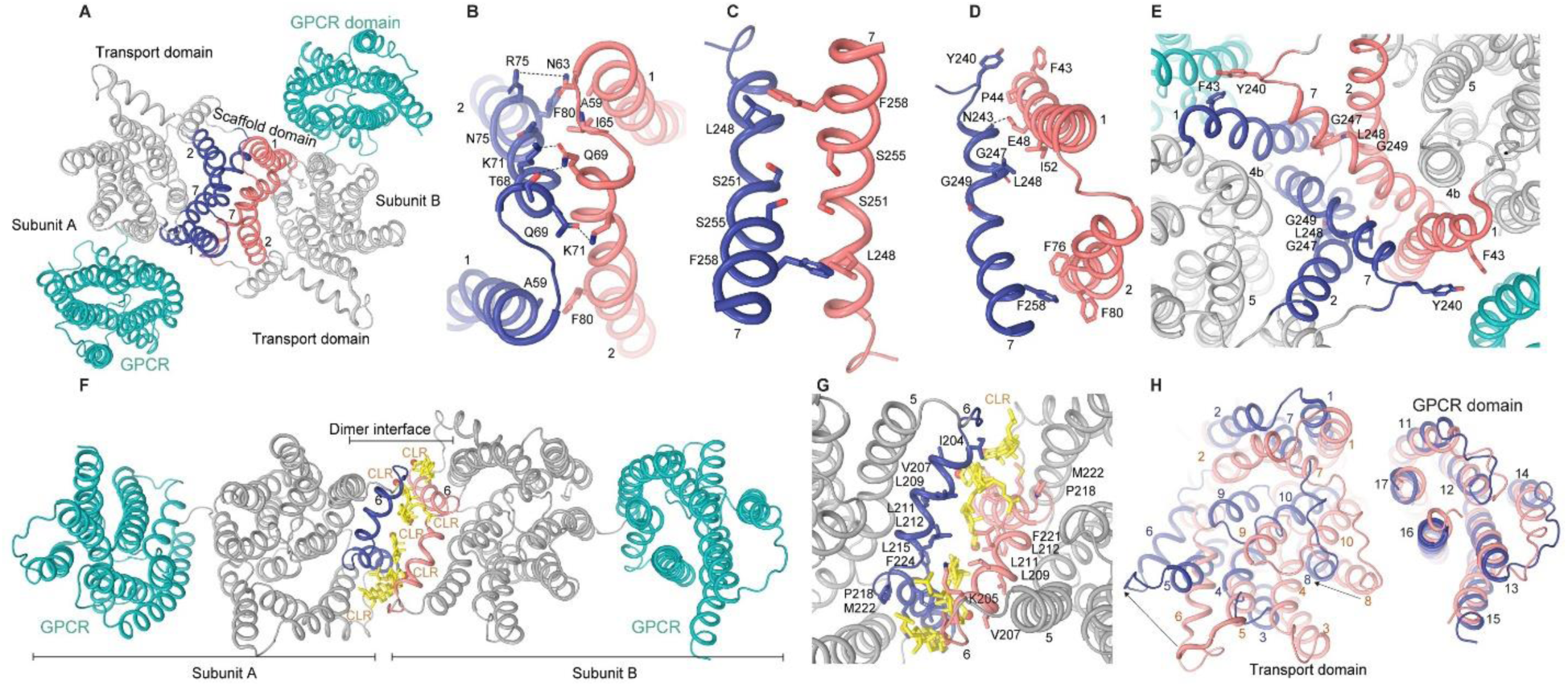
Dimeric assemblies of GPR155. **(A)** Cytoplasmic view of the dimeric interface in the PIN-like dimer, with interface helices colored salmon and blue. **(B)** Dimeric interface between TM1 and TM2, with stabilizing residues depicted as sticks. **(C)** Dimeric interface involving TM7 from each protomer, with key interacting residues shown as sticks. **(D)** Dimeric interface between TM2 and TM7, with interacting residues shown as sticks. **(E)** Luminal view of PIN-like dimeric interface. **(F)** Cholesterol-driven dimer of GPR155 with interface helices colored salmon and blue, and cholesterol shown as yellow sticks. **(G)** Enlarged view showing six cholesterol molecules that stabilize the dimer interface. **(H)** GPCR domain-based structural superimposition of monomers from PIN-like dimer and cholesterol-driven dimer.

The luminal half of the TM7 helix exhibits a bending of nearly 30° at a flexible stretch of amino acids involving the 247-GLG-249 residues, creating a unique spatial feature at the dimer interface (Fig. 4D and E). This bending results in a sizable luminal-facing cavity formed by the convergence of these bent helices (Fig. 4E and Fig. 5). The TM7 helices bending, along with the luminal halves of the TM4 and TM5 helices acting as side walls from each protomer, facilitates the fusion of two individual cavities into a single, expansive, bowl-shaped cavity measuring 30 Å × 27 Å × 14 Å (Fig. 4E and 5). The absence of other adjacent transmembrane helices flanking this cavity allows it to remain relatively open and accessible and is crucial for enabling interactions with luminal small molecules or substrates, thereby playing a significant role in the transceptor function. Two cholesterol molecules located between TM1 and TM11 in each protomer contribute indirectly to the dimeric interface. Despite the low sequence similarity, the structural features of this dimer closely mirror those found in PIN transporters, highlighting its significant structural similarity to auxin transporters (fig. S7, A-D). This evolutionarily conserved dimerization strategy across auxin transporter family protein PIN and the human GPR155 highlights shared structural and functional principles governing auxin transport in plants and lysosomal regulation in human cells.

**Figure 5.**
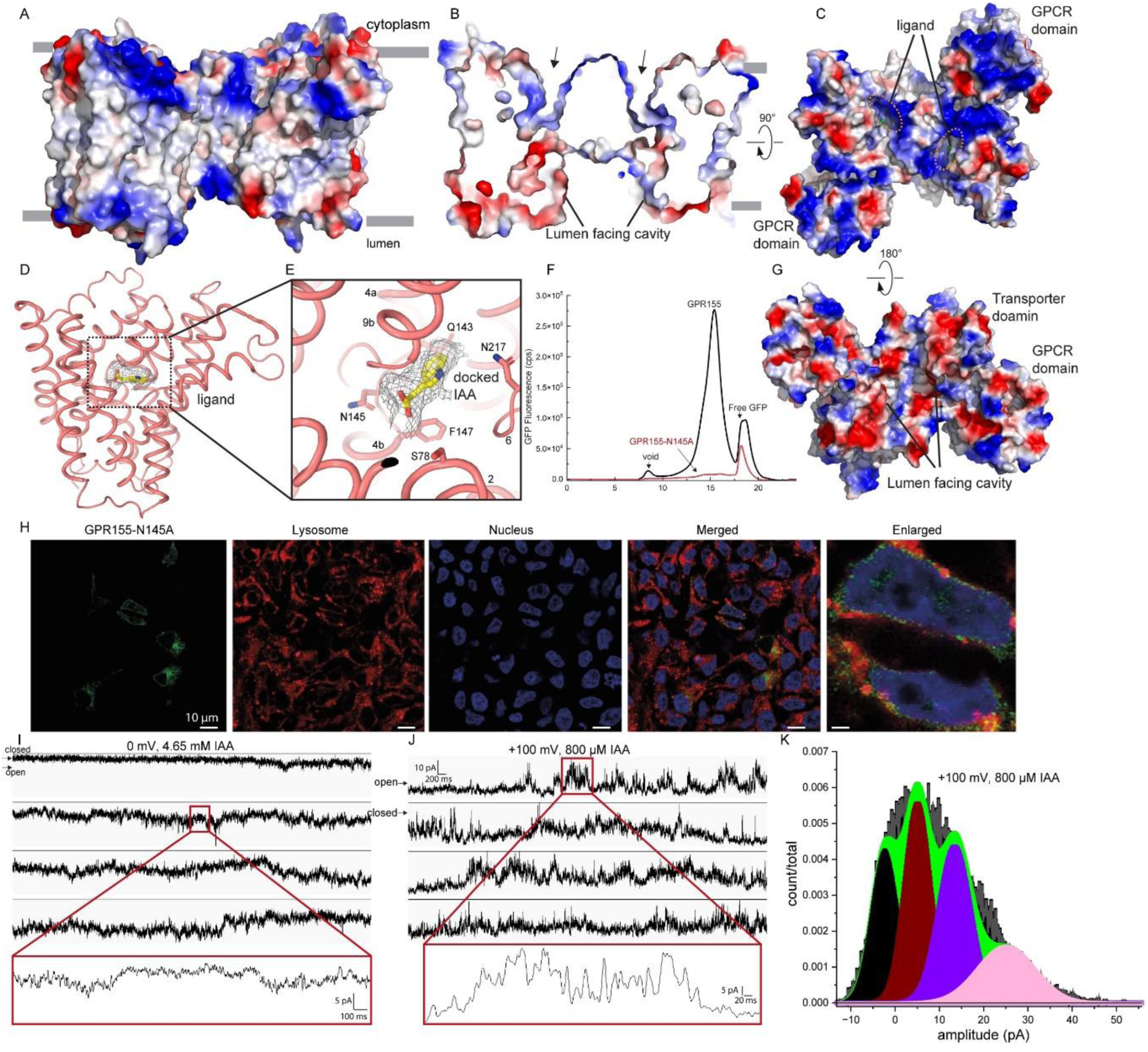
GPR155 in the inward open state. **(A)** Side view of GPR155 shown with an electrostatic surface representation. **(B)** A cutaway view of electrostatic surface representation GPR155, highlighting the cavity leading to the substrate binding pocket. **(C and G)** Cytoplasmic and luminal view of GPR155. **(D)** An unknown density in close proximity to the central X-shaped crossover, with a docked IAA molecule shown within this density as a gray mesh. **(E)** Enlarged view of the ligand-binding pocket, with residues that interact with the docked IAA depicted as sticks. **(F)** FSEC profile comparing wild-type GPR155 (dark color) and the GPR155-N145A mutant. **(H)** Confocal fluorescence images of HEK293 cells treated with LysoView^TM^ 594 for lysosome staining. **(I)** Representative lipid bilayer recordings of GPR155 reconstituted in POPC, POPG, and POPE, illustrating opening bursts at 0 mV, 4.65 mM of IAA, openings are downwards. An enlarged view of the recording is shown in the red box. **(J)** +100 mV, 800 μM, openings are upwards. An enlarged view of the recording is shown in the red box. **(K)** A histogram of amplitude distributions of transceptor currents in the trace is shown in J. The distribution was fitted with the sum of three Gaussian components considering three transceptor counts with means of 0 pA (transceptor-closed current level), 5.2 pA (1 transceptor-open current level), 13.3 pA 1 transceptor-open current level), 24.9 pA (3 transceptor-open current level).

Our second dimer structure reveals a distinct mechanism for homodimerization, where the protomers are arranged in a linear configuration, with the transporter domains sandwiched between GPCR domains on each side. This dimerization mode is primarily mediated by six cholesterol molecules embedded at the dimeric interface, and therefore we named it a cholesterol-driven dimer (Fig. 4F). These cholesterol molecules are strategically positioned between TM6 from each protomer, creating a unique cholesterol-mediated interaction that contributes significantly to the overall dimer stability (Fig. 4F and G). TM6 is characterized by a discontinuity caused by the presence of a conserved proline residue (P218) located approximately in the middle of the membrane. This proline induces a sharp bend of nearly 103°, resulting in the cytoplasmic half of TM6 lying nearly parallel to the membrane (Fig. 4G). This unique conformation allows the six cholesterol molecules to be wedged between the two TM6 helices, one from each protomer, stabilizing the cholesterol-mediated dimer. These cholesterol molecules are stabilized by an extensive set of hydrophobic interactions with hydrophobic residues, involving leucine, isoleucine, valine, and methionine from TM6 of each protomer (Fig. 4G). By embedding between the TM6 helices, they create a stable interface that supports the dimer structure without the need for direct helix-helix interactions. This arrangement of the protomers results in a length of 148.79 Å, as measured between the Q478 residues, which is significantly longer compared to the 106.6 Å measurement in the PIN-like dimer (Fig. 1, fig. S6 D, G). The primary cause of this difference lies in the distinct dimerization modes: in the PIN-like dimer, the GPCR domains are situated close to the dimeric interface, giving the overall structure a rhomboid shape. In contrast, in the cholesterol-driven dimer, the GPCR domains are positioned distally from the dimeric interface, leading to a more elongated and linear arrangement of the two monomers (Fig. 4A, 4F, and fig. S6 D-I).

The cholesterol-driven dimer lacks the phospholipid at the transporter and GPCR interface, providing an excellent template for understanding the role of phospholipids and cholesterol in transceptor and oligomeric assembly. Structural superimposition based on the GPCR domain of monomers from cholesterol-driven dimer and PIN-like dimer revealed that the luminal side of the transporter remained nearly static while the cytosolic side exhibited pronounced structural changes. Moreover, this structural analysis revealed dramatic rigid body movement in the transporter domain, with its TM6 exhibiting a movement of nearly 14.5 Å (Fig. 4H). In the cholesterol-driven dimer, the cytoplasm-facing regions of the GPCR domain and transporter domain moved apart by 5.2 Å (measured between E341 and F711) when compared to PIN-like dimer, suggesting the role of phospholipids in stabilizing the compact assembly of tranceptor and consequently PIN-like dimer assembly of GPR155. The loosening at the transceptor interface is propagated via the TM9-TM10 linker to the cytosolic half of TM6, which is moved away from the transceptor interface by 3.5A, subsequently leading to the formation of the cholesterol-driven dimer.

### GPR155 in inward-open state

The surface charge analysis of GPR155 revealed a pronounced positively charged region on the cytoplasmic side of the transceptor, while the luminal side exhibited a relatively negative charge, suggesting a potential role for charge in substrate recognition and transport directionality (Fig. 5A). Additionally, GPR155 features a solvent-accessible cavity with a weak positive potential that extends from the cytoplasmic side to the central X-crossover region, indicative of an inward-open state of the transporter domain (Fig. 5 B and C). Indeed, GPR155 shows a more substantial alignment with the inward-facing structures of inward-open conformation observed in the NPA-bound form of the PIN8 transporter (fig. S7 A-D, H).

A small non-protein map density was detected within the positively charged substrate-binding pocket, located near the central X-shaped crossover. This density partially overlaps with the bound NPA in the PIN8 structure, suggesting that it could represent an endogenous ligand carried over protein purification. Interestingly, this density can be appropriately docked with auxin and auxin-like molecules (Fig. 5D, E) (*3*). The positioning of this non-protein density suggests interactions with key ligand binding residues: Q143, N145, and F147 from TM4, as well as S78 from TM2 and N217 from TM6. Specifically, the carboxyl group of docked IAA appears to form a hydrogen bond with N145 in TM4, while the imidazole ring engages in hydrophobic interactions with F147 in TM4, F332 in TM9, and I219 in TM6. Interestingly the key residue N145, involved in IAA binding, is evolutionarily conserved between PINs and GPR155, suggesting its critical role in substrate binding and a conserved mechanism of substrate transport across these transporters (Fig. 5D and E). Substituting N145 with alanine in GPR155, resulted in a strong phenotype characterized by significantly reduced expression and altered targeting to the lysosome, suggesting compromised functionality in the mutant (Fig. 5F, H). Indeed, the FSEC profile of this mutant exhibited significantly reduced expression and oligomeric assembly compared to the wild-type protein, suggesting that this residue plays an important structural and functional role. These observations highlight the structural basis for substrate binding and specificity in the GPR155 transceptor, providing insights into its functional mechanisms.

As GPR155 exhibits substantial structural similarity to the auxin transporter PIN family protein, including a conserved asparagine at the central crossover, next we investigated its auxin transport activity using lipid bilayer experiments. Upon reconstitution in a lipid mixture of POPC, POPG, and POPE, GPR155 showed concentration-dependent transport activity, as evidenced by inward currents with amplitudes ranging from approximately 5.5 pA to 20.4 pA in response to 4.65 mM IAA at zero voltage (Fig. 5 I-K). Notably, robust inward currents were not observed at lower IAA concentrations at zero voltage. These lipid bilayer experiments further revealed that voltage could enhance the IAA transport activity of GPR155. Outward currents ranging from approximately 5 pA to 30 pA at +100 mV were observed in response to 800 μM IAA, whereas the outward current amplitudes decreased to 5.5 pA with 200 μM IAA at the same positive voltage (Fig. 5I-K, fig. S10). These findings suggest that both the concentration of IAA and the applied voltage across the membrane influence the IAA transport activity of GPR155. Additionally, it is possible that other indole-containing compounds similar to IAA could also serve as substrates for GPR155. However, further research is required to elucidate the precise mechanisms by which voltage facilitates the transport function of GPR155.

## Discussion

GPR155, also known as LYCHOS, is a lysosomal membrane transceptor that integrates structural features of both transporters and G protein-coupled receptors (GPCR) (*12*). This transceptor binds to cholesterol and regulates lysosomal signaling pathways. In this work, we present structural insights into GPR155, revealing its molecular architecture, potential transport mechanism, and role of cholesterol in stabilizing distinct assemblies. The interface between the transporter and GPCR domains is stabilized by a pair of phospholipids, similar to the compact transmembrane dimerization observed in GPR158 (*26, 27*). These phospholipids effectively staple two different classes of membrane proteins together, potentially offering a novel mechanism for transceptor modulation.

The transporter domain of GPR155 exhibits structural similarity to the auxin transporter PIN family of proteins within the BART superfamily (*35*). Despite low sequence similarity, GPR155 adopts the NhaA fold and dimerizes similarly to auxin transporters, revealing an overlooked structural link between plant auxin transporters and their mammalian counterparts. This exemplifies convergent evolution, where GPR155, despite its distinct origins, has developed transport mechanisms reminiscent of plant auxin transporters, underscoring the evolutionary parallels in cellular transport systems. The transporter domain in all the structures presented here is captured in an inward-open state, which closely resembles the inward conformations of the auxin transporters PIN1, PIN3, and PIN8 (*3–5*). In these structures, we observed a small non-protein density at the weakly positive X-shaped crossover support site, suggesting it may bind negatively charged substrates like auxin or auxin-like molecules. Indeed, our functional studies, using lipid bilayer experiments validate the IAA transport activity of GPR155. Although we were unable to capture the outward-open state of the transporter domain, its structural comparisons with the PIN family proteins suggest an evolutionarily conserved elevator-type mechanism of substrate transport and regulation.

Structural-based sequence analysis of the GPCR domain of GPR155 classifies it as a Class B receptor based on conserved motifs in TM4 and TM7 of classical GPCRs. Interestingly, LL7 of GPR155 adopts a unique fold that plugs the orthosteric site, resembling an agonist-like conformation seen in Class A GPCRs (*42–44*). As GPR155 localizes to lysosomes, it is less likely to be activated by an extracellular ligand, instead, the unique conformation of LL7 may keep the receptor in an activated state upon engagement of the GPCR domain with cytosolic proteins.

Further, our study reveals dynamic regulation of GPR155 oligomeric states by cholesterol and phospholipids, uncovering two distinct dimerization modes. PIN-like dimer raises the possibility of a potential link between GPCR activation and auxin transport in lysosomes (*3–5*). Cholesterol has been previously proposed as an endogenous ligand of GPR155, (*12*), however, our structural analysis reveals multiple cholesterol binding sites on GPR155, including those on both the GPCR and transporter domains, indicating a broader role for cholesterol in defining different dimerization assemblies and regulation of GPR155. Further studies are needed to test these mechanisms. The interplay between auxin and cholesterol opens new avenues for future research into the role of GPR155 in lysosomal biology and its broader implications in cellular signaling pathways.

A key question arises: does the GPCR domain influence substrate transport within the transporter domain of GPR155? The fusion of this GPCR domain to the transporter domain, along with its specific lysosomal expression, hints at a unique activation mechanism for this C-terminus 7TM bundle, potentially independent of extracellular ligands. Indeed, we observed LL7 loop occluding the orthosteric ligand binding pocket which could potentially act as an activator for the class B GPCR domain. This GPCR domain features a LYCHOS effector domain (LED) within an exceptionally long ICL8 loop, spanning nearly 110 residues. Although disordered in our current structure, this loop may become ordered upon interaction with GATOR1 complexes. Furthermore, the C-terminus of the GPCR culminates in a Disheveled, Egl-10, and Pleckstrin (DEP) domain, commonly found in the RGS family of proteins, which interact with Gβ5-γ subunits (*26, 27*). This raises intriguing possibilities for the function of GPR155 —could it bind Gβγ subunits? Exploring these hypotheses will be crucial in understanding the full scope of functional implications of GPR155 in auxin/cholesterol-based cellular signaling and lysosomal function. This work lays a solid foundation for future research to further investigate the role of GPR155 in lysosomal biology.

## Author Contributions

MS, DKJ, SS, and AKS designed the project, built models, and analyzed data. MS and DKJ designed constructs, developed expression and purification protocols, prepared protein samples, and carried out cryo-EM data collection and processing. MS carried out FSEC and SEC experiments with help from SN MS and SS built the models of GPR155. MS and AKS carried out lipid bilayer recording. SS and AKS supervised the project. MS, DKJ, SS and AKS wrote the manuscript with inputs from other authors.

## ACKNOWLEDGEMENTS

We thank the staffs Adam Wier, Thomas Edwards, and Tara L. Fox, of the NCI National Cryo-EM Facility for data collection. The authors thank the Sophisticated Analytical & Technical Help Institutes (SATHI) Foundation, IIT Delhi, and Indian Science Technology and Engineering Facilities Map (I-STEM), National Cryo-EM Facility, and Sandeep Singh for sample screening and data collection.

## Data availability

The cryoEM density maps and Coordinates have been deposited in the Electron Microscopy Data bank (EMDB) and Protein Data Bank (PDB), respectively, with accession codes XXXX, XXXX.

## Materials and Methods

### Constructs design

We synthesized a gene that encodes human GPR155 (residues 1-870) for expression in mammalian cells. The Gene Universal (Newark) synthesized this gene, which was inserted into a modified pEG BacMam vector. Following the C-terminus of GPR155 in this vector, there is a thrombin cleavage site (LVPRG), an enhanced green fluorescent protein (eGFP) tag, and a Strep-tag (WSHPQFEK) for fluorescence size-exclusion chromatography (FSEC) screening (*46*). Additionally, a version of the construct without the eGFP tag was prepared for large-scale protein production. Standard molecular biology techniques, including restriction enzyme digestion and ligation, were employed to generate its variants.

### GPR155 expression and purification

We generated and amplified recombinant bacmids and baculovirus encoding GPR155 in Sf9 insect cells (ThermoFisher Scientific, Cat. No. 11496015) following established protocols (*46*). To produce the baculovirus, Sf9 cells were transfected with approximately 21 μg of bacmid DNA using Cellfectin. After an incubation period of 5-6 days, the resulting virus was harvested, filtered through a 0.22-micron filter, and stored at 4°C. This baculovirus was then used to infect 600 ml of Sf9 cells cultured in SF900^TM^ III media (Thermo Fisher Scientific, GIBCO #12659017) for 88-96 hours, followed by a 20-fold concentration of P2 baculovirus at 63900 g using T-647.5 rotor (Thermo Fisher) and Sorvall WX 100 Ultra Series Centrifuge. The GPR155 was expressed in suspension-adapted HEK293F cells (Thermo Fisher Scientific, GIBCO Cat. No. R79007) that were cultured at 37°C with 5% CO2 in FreeStyle 293 Expression Medium (Gibco-Life Technologies #12338-018). To produce the GPR155 protein, we adapted protocols with slight modifications from those established for TRPV6 (*47*) and GPR158 (*26*). HEK293F cells were cultured at 37°C to 2.3-2.8 × 10^6^ cells/ml before being infected with P2 BacMam viruses of GPR155. 8-10 hours post-infection, GPR155 expression was boosted by adding 10 mM sodium butyrate. The cultures were incubated for an additional 61-63 hours at 30°C before the cells were harvested by centrifugation at 5,422g for 20 minutes using a rotor (F9-6×1000 LEX) in a Sorvall LYNX 6000 centrifuge (Thermo Fisher Scientific). The harvested cell pellet was washed in PBS (pH 8.0) and pelleted by spinning for 15 minutes using an A27-8×50 rotor and Sorvall LYNX 6000 centrifuge. All the steps of GPR155 protein purification were conducted at 4 °C. For GPR155 purification, the cell pellet was resuspended lysis buffer containing 20 mM Tris-Cl pH 8.0, 150 mM NaCl, 1.50 mM PMSF, 4.50 μM Leupeptin, 1.50 μM Pepstatin A, 1.0 μM Aprotinin,), followed by lysing using Sonics Vibra-Cell VCX 750 Sonicator with 12×3 cycles of 10 seconds “on” at amplitude 20, followed by 15 seconds “off,” while keeping the sample on ice to prevent heat denaturation. Following cell disruption, cell debris was removed by centrifugation of lysate at 9,900g for 18 minutes at 4°C. The resulting supernatant was subjected to solubilization in a solution containing 1% (w/v) lauryl maltose neopentyl glycol (LMNG) supplemented with 0.1% (w/v) cholesteryl hemisuccinate (CHS) while stirring for 1 hour and thirty minutes at 4°C. After solubilization, the mixture was ultracentrifuged at 1,77,500g and 4°C for 1 hour to remove insoluble debris using T-647.5 rotor (Thermo Fisher) and Sorvall WX 100 Ultra Series Centrifuge. The resulting clear soluble fraction was allowed to bind to anti-STREP resin for 12-17 hours at 4°C. The non-specific bound proteins were removed from the STREP resin by passing 20-30 column volumes of size-exclusion chromatography (SEC) buffer (20 mM Tris-HCl (pH 8.0), 150 mM NaCl, 0.01% LMNG, and 0.001% CHS) through the Econo-Pac gravity chromatography column (BIO-RAD). Following the washing of STREP resin, 10 ml of elution buffer, which is the same as SEC buffer but with 2.5 mM D-desthiobiotin, was passed through the gravity column to elute bound GPR155 protein. The affinity-purified GPR155 protein was concentrated to 0.51 ml using a 100 kDa MWCO amicon ultra centrifugal filter (Millipore). The concentrated protein solution was centrifuged at 66,000g and 4°C for 30 minutes using a tabletop Micro-Ultracentrifuge (Sorvall MTX 150). The clarified sample was injected into a Superose™ 6 10/300 GL size-exclusion chromatography (SEC) column connected to an ÄKTA^TM^ pure system, pre-equilibrated with SEC buffer. The peak fractions corresponding to the GPR155 peak from the SEC column were collected and concentrated to 0.72 mg/ml for structural studies using cryo-EM. For lipid bilayer recordings, the GPR155 protein was purified in LMNG detergent without CHS using the same purification protocol.

### Cryo-EM sample preparation and data acquisition

Au/Au grids were prepared using the previously developed protocols (*48*). Briefly, C-flat CF-1.2/1.3-2Au holey carbon grids, purchased from Protochips, Inc, were gold-coated with ∼60 nm layer of gold using an Edwards Auto 306 evaporator. The gold-coated grids were then with Ar/O_2_ (4 minutes, 50 watts) using a Gatan Solarus (model 950) to remove the carbon. Prior to sample application, the grids underwent a final H_2_/O_2_ plasma treatment (25 seconds, 10 watts) to make the surfaces hydrophilic. The Cryo-EM grids for imaging were prepared by applying 3μL of purified GPR155 protein on the gold side of grids held inside a FEI Mark IV Vitrobot (Thermo Fischer Scientific) with temperature set at 4°C and humidity maintained at 100%. A blot force of 2.5, blot time of 2.5 seconds, and wait time of 20 seconds were used for the blotting process. The excess liquid from the grid was removed during the blotting process and was then plunge-frozen in liquid ethane. The single particle images of the frozen-hydrated sample of GPR155 were acquired on a 300 kV Titan Krios transmission electron microscope (TEM) equipped with a Gatan K3 Summit direct electron detection (DED) camera (Gatan, Pleasanton, CA, USA) and a post-column GIF Quantum energy filter operating in counting mode at a magnification of 81,000, resulting in a pixel size of 1.09 Å. A total of 13,970 movies were acquired with a defocus range of −1.6 to −2.5 μm. Data acquisition was done at a total dose of 50 e−Å^2^ across 40 frames with a total exposure time of 2.6-s giving a dose rate of ∼22.71 e−/s/phys pixel per frame.

### Image processing and 3D reconstruction

The dataset was processed using RELION(*49*) and CryoSPARC (*50*), or both (refer to figs. S3, S4, S5, and table S1 for details). The 40 frames in each movie were corrected for beam-induced motion using patch motion correction, followed by the estimation of the contrast transfer function (CTF) using CryoSPARC with the patch CTF estimation tool (*50*). Following CTF estimation, the exposure curation tool available in CryoSPARC (*50*) was used to reject poor quality micrographs, viz those with a CTF fit value lower than 6 Å, in addition to outliers astigmatism, resulting in 11,572 micrographs for further processing. A small subset of micrographs was used initially to select representative particles/templates using blob and template picker, in CryoSPARC, followed by reference-free 2D classification. Finally, a TOPAZ model was first trained against a good set of ∼240,000 particles encompassing all three classes of GPR155 and then this trained model was used for particle extraction on the entire dataset (*51*). The TOPAZ extraction produced a total of 6,757,838 particles. Following extraction, the particles were aligned using multiple rounds two-dimensional class averaging and subsequently removing the junk particles; this was followed by ab-initio reconstruction, heterogenous, homogenous, and non-uniform (NU) refinements, resulting in 151,034 particles for monomer, 135,237 particles for PIN-like dimer, a subset of 123,102 particles for cholesterol-driven elongated dimer. Because of the modular design of GPR155, without any discernible density corresponding to the luminal or cytoplasmic domain, the TM region was not well resolved in our structures. Thus, focused and local refinements with a mask around the TM region or the transporter/GPCR domains were used to further improve the density in these regions. The reported resolution of the final maps was estimated with the gold-standard Fourier shell correlation (GSFSC) at FSC=0.143 cut-off as implemented in CryoSPARC. The local resolution estimation was done using the local resolution estimation tool in CryoSPARC (*50*). EM density of the reconstructed maps was visualized using UCSF Chimera(*52*).

### Model building and refinement

The cryo-EM reconstruction of the PIN-like dimer of GPR155 achieved an overall resolution of 3.45 Å with the transmembrane (TM) domain resolved better compared to monomeric and cholesterol-driven dimeric reconstructions. We utilized a model predicted by AlphaFold as a reference to build the TM domain of the PIN-like dimer of GPR155 using real-space refinement in COOT (*53*). The side chain density corresponding to bulky amino acids such as tryptophan, tyrosine, phenylalanine, and arginine helped in correct sequence assignment, with each residue manually adjusted and iteratively built into the cryo-EM density using COOT (*53*). The model of the monomeric subunit resulting from the PIN-LIKE dimer was then docked in cryo-EM maps of monomeric and cholesterol-driven dimer of GPR155 using UCSF Chimera (*52*) to build corresponding models in the COOT (*53*). All models underwent real-space refinement against their respective cryo-EM maps with restrained group ADP refinement and were subsequently validated using comprehensive cryo-EM validation in PHENIX (*54*). The UCSF Chimera (*52*), ChimeraX (*55, 56*), and PyMOL (*57*) were for used structure visualization and figure preparation. The cryo-EM data collection parameters and refinement statistics are shown in Table S1.

### Planar lipid-bilayer recordings

To assess the transporter activity of the purified GPR155 protein, we conducted planar lipid bilayer experiments following previously described protocols (*58*). First, we prepared a lipid mixture consisting of 1-palmitoyl-2-oleoyl-glycero-3-phosphocholine (POPC), 1-palmitoyl-2-oleoyl-glycero-3-phosphoethanolamine (POPE), and 1-palmitoyl-2-oleoyl-glycero-3-phosphoglycine (POPG) at a 3:1:1 ratio in a 30 mM solution dissolved in n-decane. This solution was used to form a bilayer across an aperture of 150 μm diameter in a Meca chip supplied by Nanion, equipped with integrated Ag/AgCl microelectrodes. The bilayer separated symmetric bathing solutions containing either 10 mM HEPES, pH 7.4, or 117.5 mM KCl and 10 mM HEPES, 1 mM MgCl2, pH 7.4. The resulting bilayer had a capacitance range of 5 to 15 picofarads. To incorporate the GPR155 protein, a micellar solution of GPR155 at a concentration of 0.05 μg/ml was mixed with 10 μM indole-3-acetic acid (IAA) and incubated with an equal volume of the lipid mixture at 30°C for 30 minutes before painting the bilayer using a brush. For zero-voltage data collection, the protein was not incubated with IAA before painting the bilayer; instead, IAA was applied to the cis compartment of the Meca chips, creating an asymmetric solution for concentration-dependent measurements of GPR155 activity. The addition of IAA to the cis compartment resulted in measurable currents, confirming the successful incorporation of the transporter protein into the bilayer and its activity. For voltage-dependent measurements of GPR155 transport activity, a symmetric solution with the appropriate IAA concentration was used. The integrity of the bilayer was regularly checked by measuring its capacitance, and baseline measurements were recorded to identify changes attributable to transporter activity. The data was analysed using software provided with nanion instrument and QUB (*59, 60*) software to determine the transport activity as well as the conductance properties of the GPR155 protein (*61*).

### Confocal fluorescence imaging

For the confocal fluorescence imaging experiments, we first, coated the circular coverslips (0.17mm thick) with 0.1% gelatin for 25 minutes at room temperature (RT) and washed them three times with 1X PBS. HEK 293 cells (CRL-1573^TM^, ATCC) were cultured on the gelatin-coated circular coverslips and were transfected with C-terminus GFP-fused GPR155 and its variant. After 48 hours of transfection, the cells were incubated with LysoView^TM^ 594 (Biotium) at a 1:1000 dilution in cell growth media for 30 min and then fixed in 4% paraformaldehyde for 12 min at room temperature. Images were captured using a fluorescence microscope (Zeiss) with a 100X objective with immersion oil, and further analysis was done using Image J software (*62*).

---

**Fig. S1.**
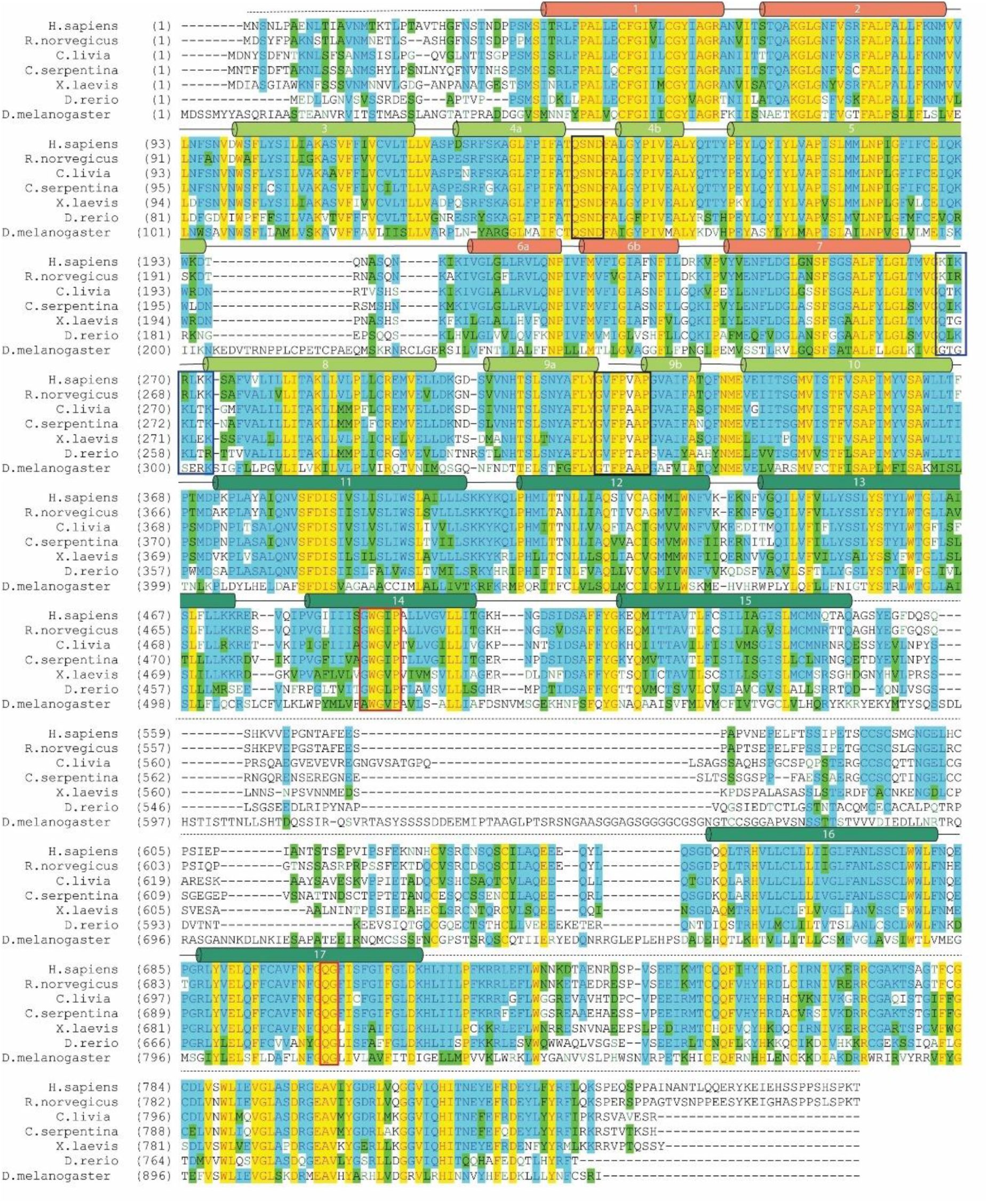
Sequence alignment of GPR155 homologs. The conserved motifs of class B GPCR are shown in the red box, while the residues interacting with the lipids are shown in the blue box. The residues that form an X-shaped crossover are displayed in the black box. In D. melanogaster, GPR155 homolog is an anchor protein.

**Fig. S2.**
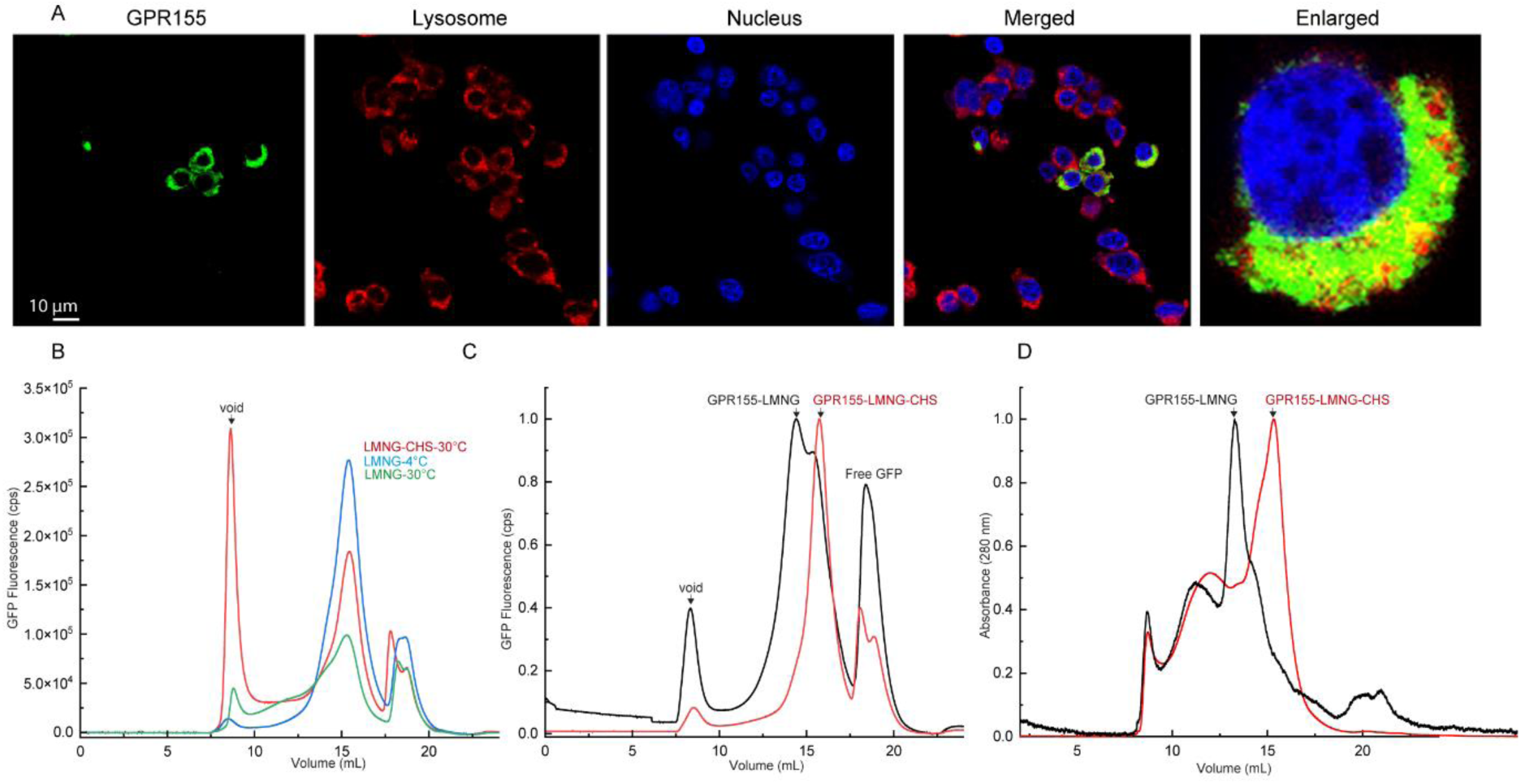
GPR155 localization and FSEC analysis. **(A)** Confocal fluorescence imaging showing the lysosomal localization of GPR155. **(B)** Thermostability analysis of GFP-labeled GPR155 protein, with and without cholesterol (CHS). **(C)** Normalized FSEC plots of GPR155 protein solubilized in LMNG detergent in the presence and absence of CHS. **(D)** Normalized size exclusion chromatography profile of GPR155 was conducted with LMNG and LMNG-CHS.

**Fig. S3.**
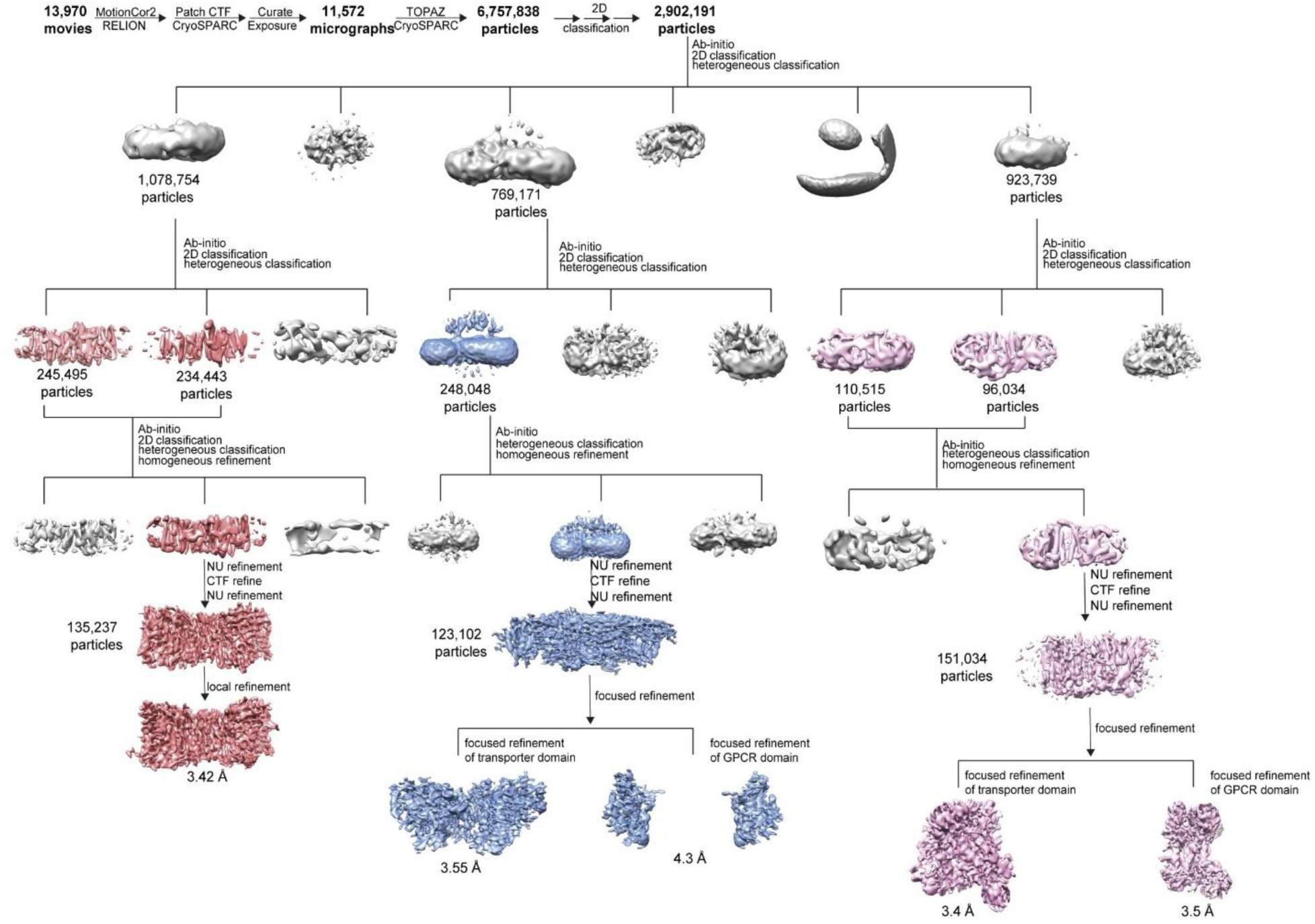
Cryo-EM processing workflow of GPR155.

**Fig. S4.**
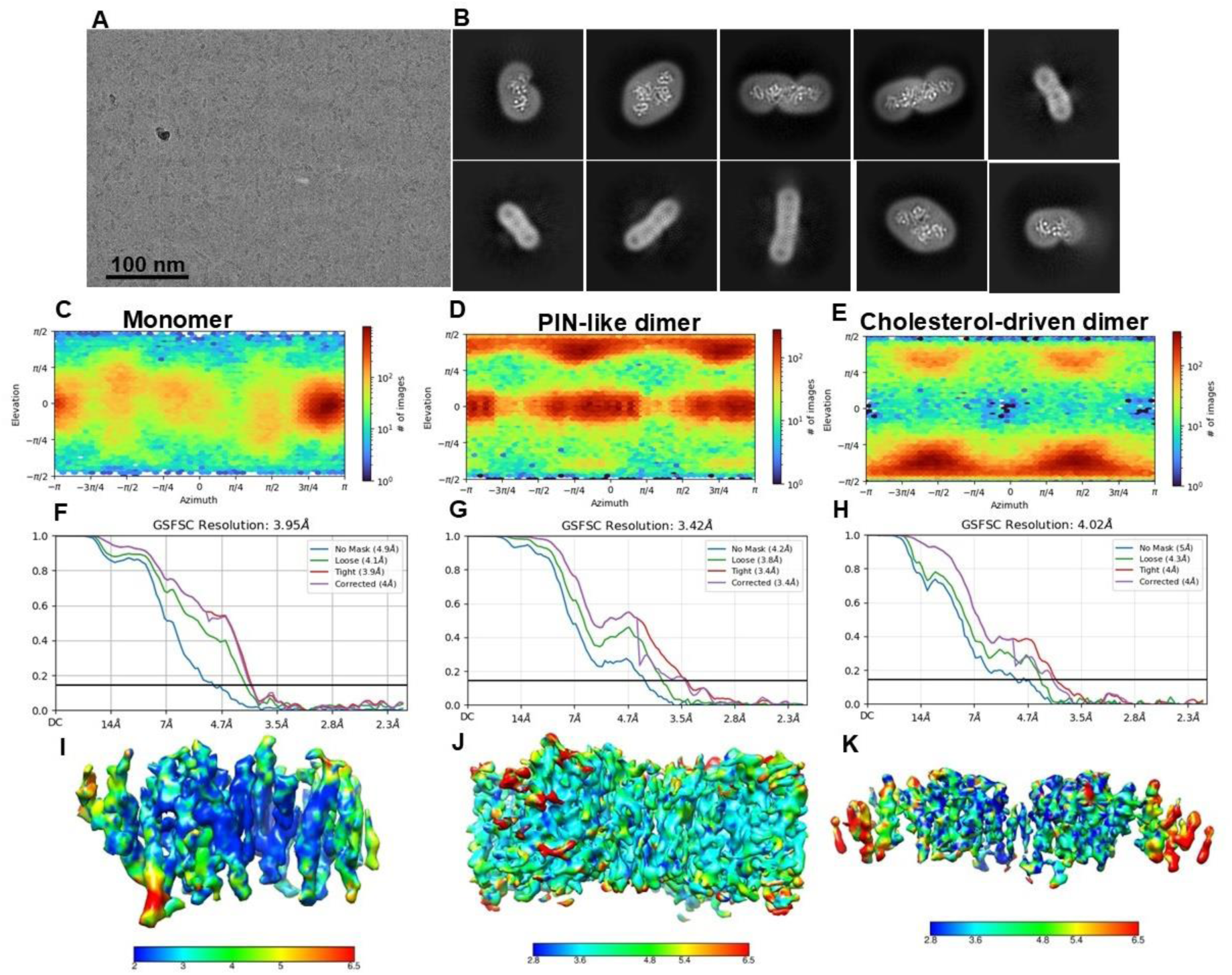
cryo-EM processing of GPR155. **(A)** Exemplary cryo-EM micrograph showing GPR155 particle in different views. **(B)** 2D class averages showing different views of GPR155. **(C, D, E)** Distribution orientation of GPR155 assemblies. **(F, G, H)** FSC curve for resolution estimation. **(I, J, K)** local resolution map of GPR155 assemblies.

**Fig. S5.**
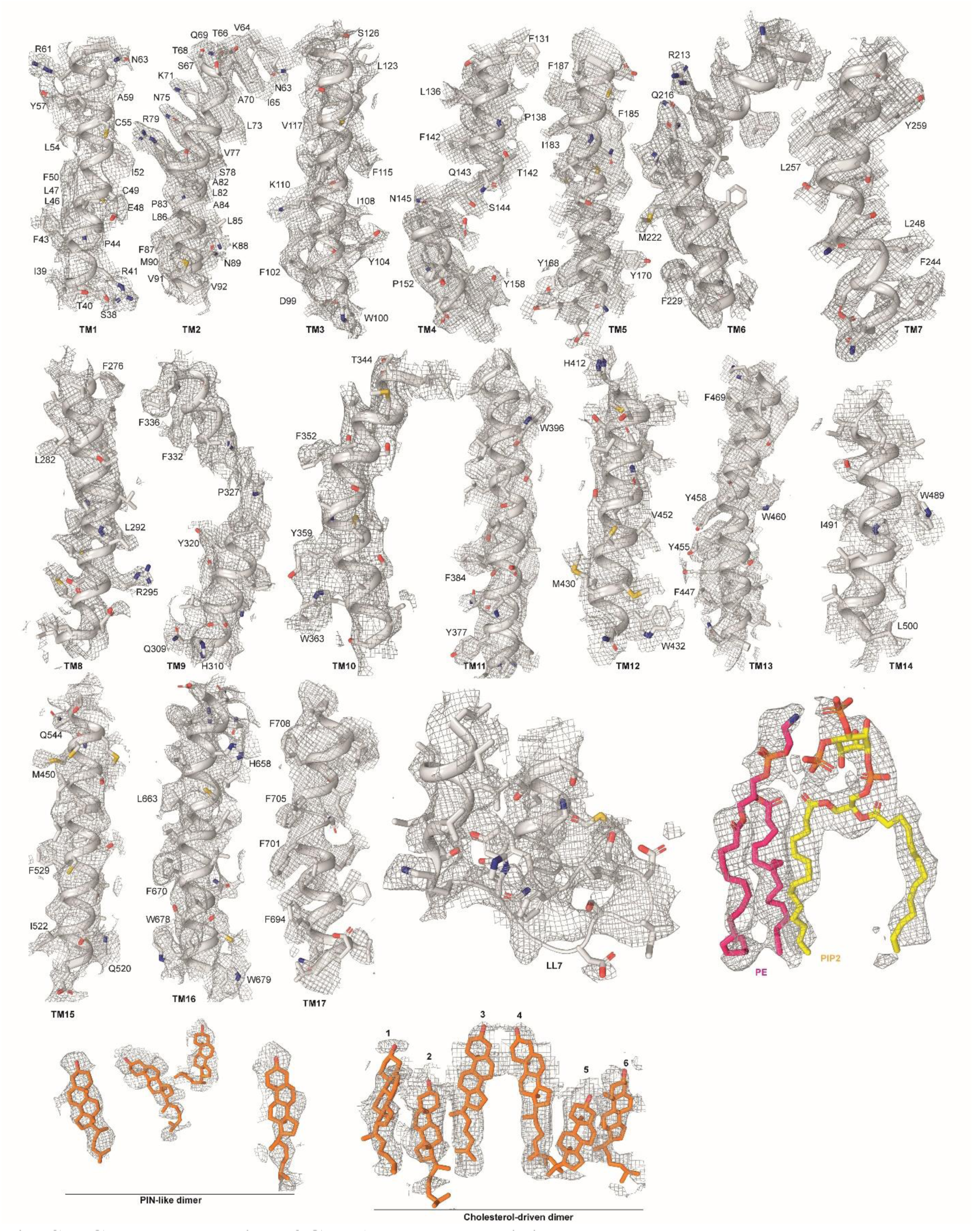
Cryo-EM density of GPR155 TMs and lipids.

**Fig. S6.**
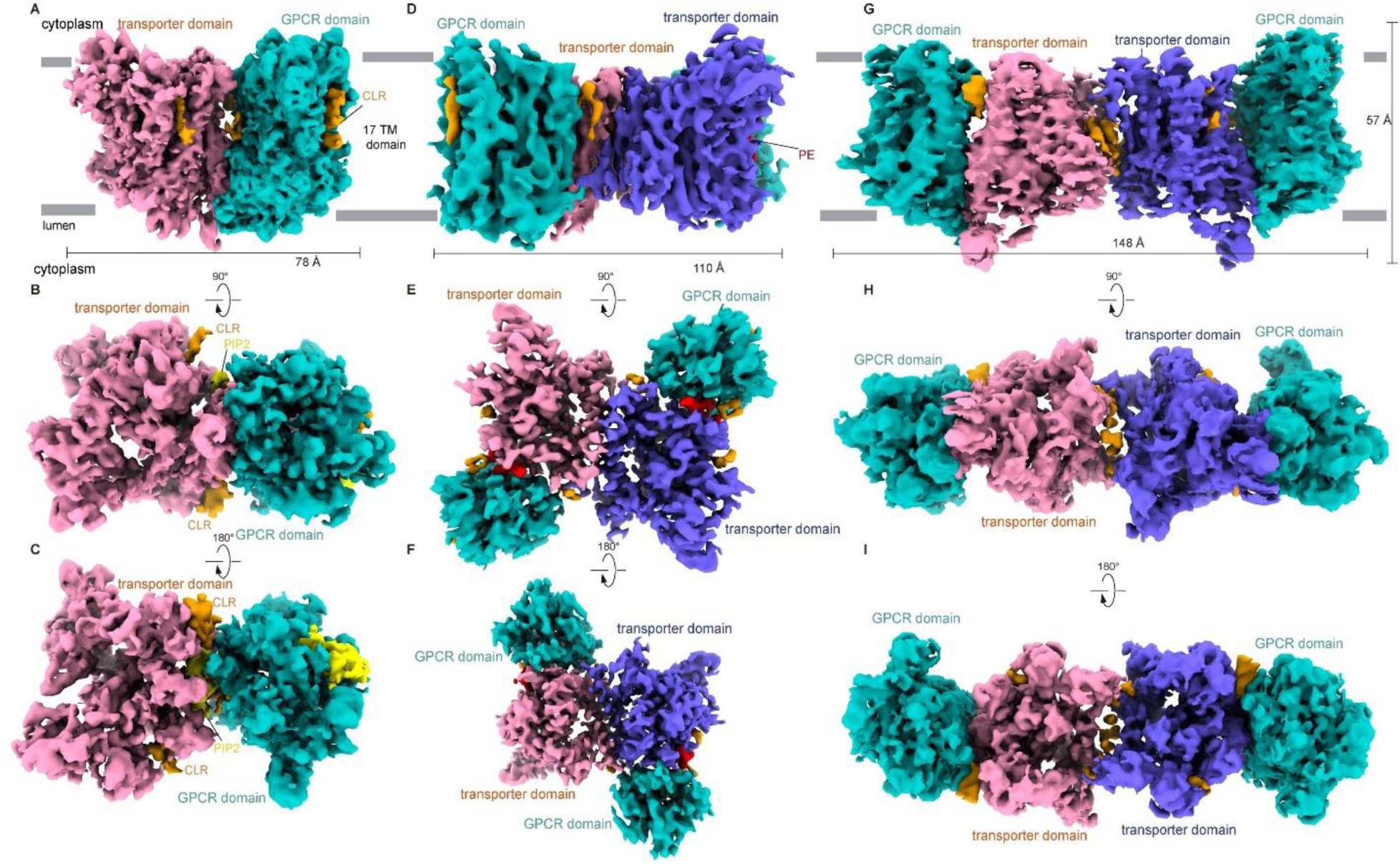
Overall Structure and Assemblies of GPR155. Cryo-EM maps of GPR155 are shown in different states: **(A)** Monomeric form, with the transporter domain (salmon) and GPCR domain (teal) viewed from the side. **(B)** Luminal view of the monomeric form. **(C)** Cytoplasmic view of the monomeric form. **(D)** PIN-like dimeric form viewed from the side. Its transporter domains in PIN-like dimer are shown in purple and salmon colors, and GPCR domains are shown in teal color. **(E)** Luminal view of the PIN-like dimer. **(F)** Cytoplasmic view of the PIN-like dimer. **(G)** Cholesterol-driven dimer, with domains colored as in D, viewed from the side. **(H)** Luminal view of the cholesterol-driven dimer. **(I)** Cytoplasmic view of the cholesterol-driven dimer. Intrinsically bound lipids are shown as surfaces: PIP2 (yellow), PE (red), and cholesterol (orange). The membrane region is marked in gray, with the cytoplasmic side on top and the lumen on the bottom.

**Fig. S7.**
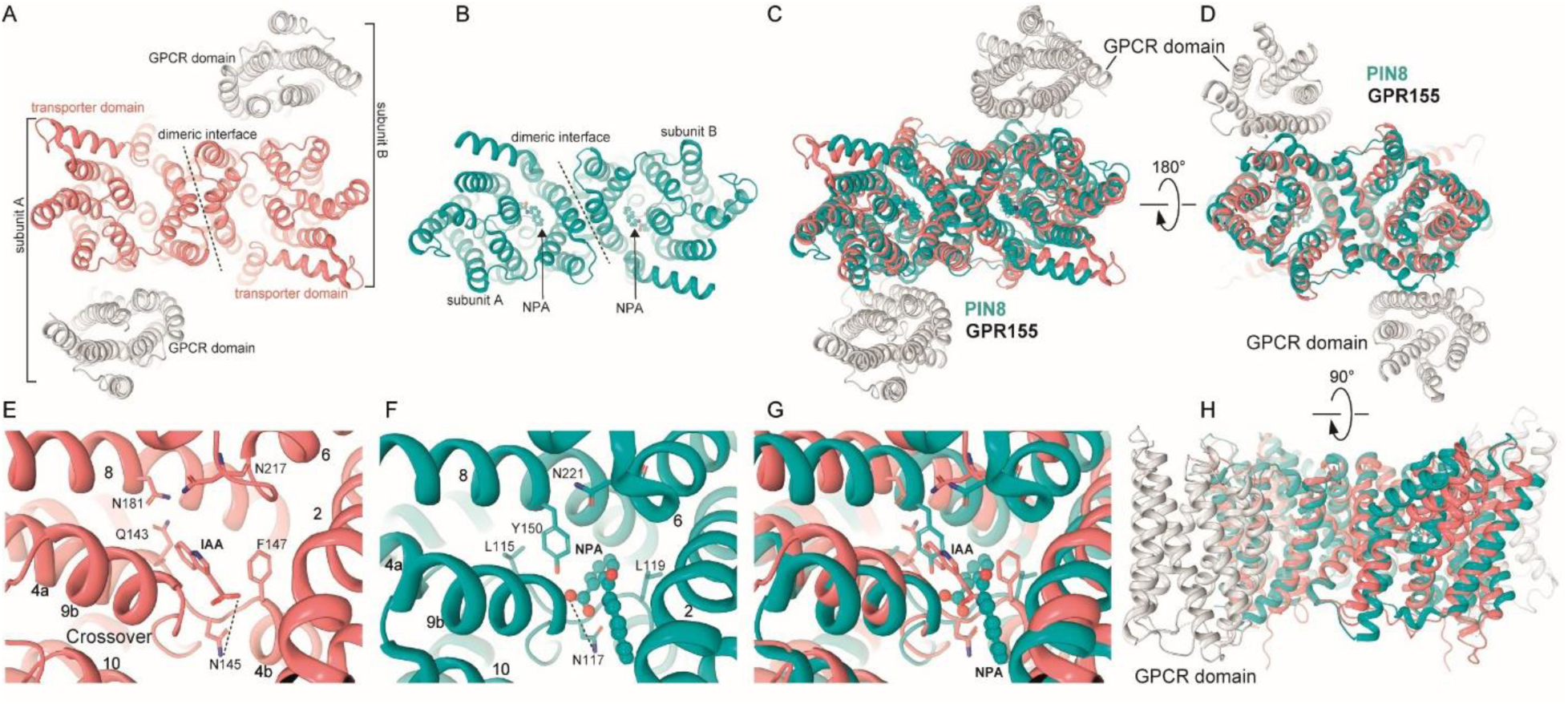
Structural Comparison of GPR155 and PINs. **(A)** Cytoplasmic view of GPR155 with transporter domain shown in salmon color, GPCR domain in gray color, and the dimeric interface is lined by a dashed line. **(B)** Cytoplasmic view of PIN8 (PDB: 7QPC) with each protomer shown in cartoon in teal color. **(C)** A view of structural superimposition of PIN8 and GPR155 from the cytoplasmic side. **(D)** A view of structural superimposition of PIN8 and GPR155 from the luminal side. **(E)** Close-up view of the ligand-binding pocket of GPR155, showing the density-mediated docked IAA molecule interacting with N145 and F147 from the crossover. **(F)** Close-up view of the ligand-binding pocket of PIN8, showing the NPA inhibitor-bound molecule interacting with N117, L119, L115, and Y150. **(G)** Structural superimposition of the ligand-binding pockets of PIN8 and GPR155, demonstrating that the density-mediated docked IAA occupies a similar position to NPA in PIN8. **(H)** A side view of structural superimposition of PIN8 and GPR155.

**Fig. S8.**
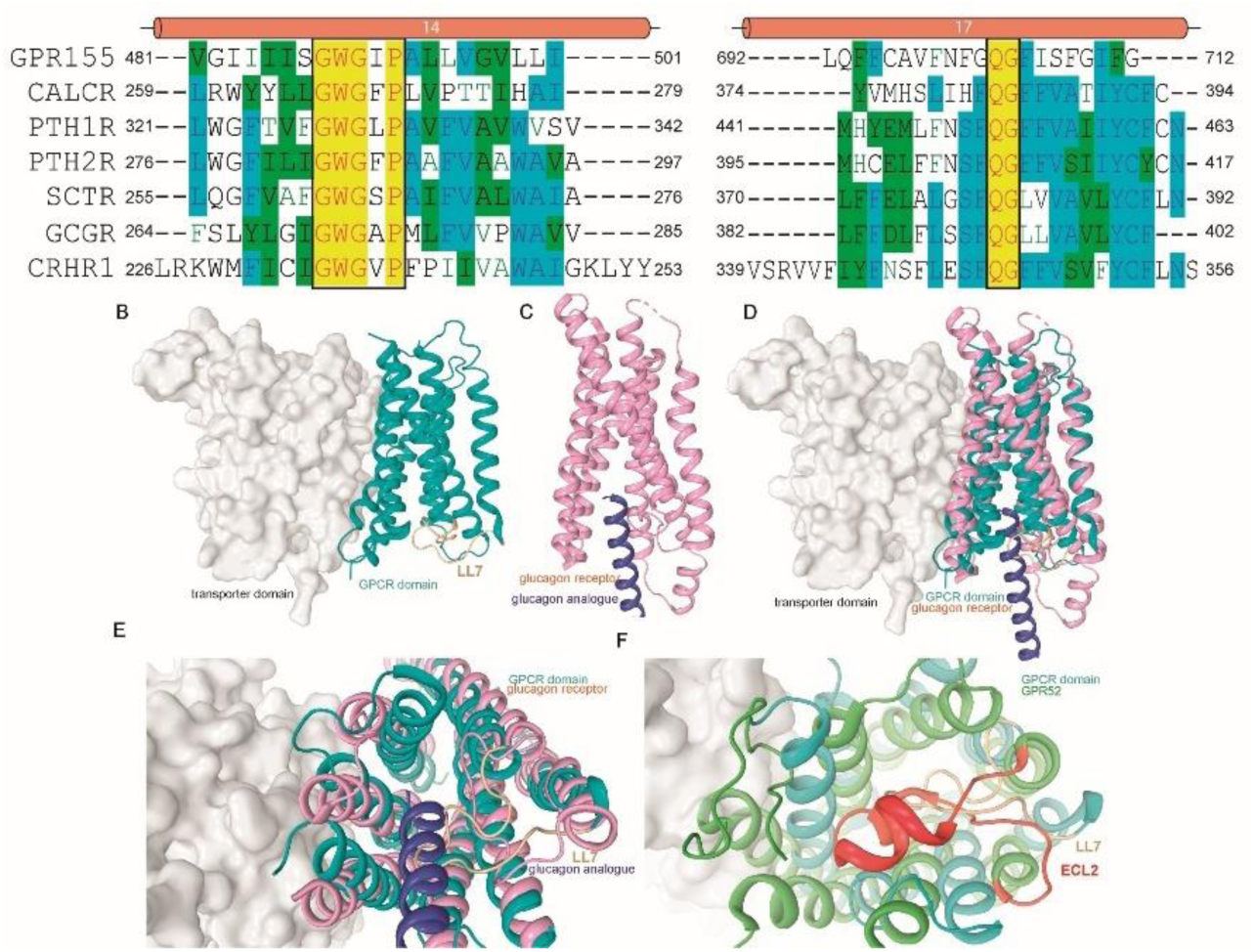
Structural comparison of GPR155 with Class B and A GPCRs. **(A)** Sequence alignment of TM14 and TM17 of GPR155 with class B GPCRs, highlighting conserved motifs in the black box. **(B)** Side view of GPR155, with the transporter domain shown as a gray surface, the GPCR domain (teal), and the LL7 loop (wheat color) represented in the cartoon. **(C)** Side view of the class B GPCR glucagon receptor (pink) cartoon representation, with the bound glucagon peptide shown in blue (PDB: 5YQZ). **(D)** Structural superimposition of GPR155 with the glucagon receptor. **(E)** Luminal view of the superimposed GPR155 and glucagon receptor. **(F)** Structural superimposition of GPR155 with Class A GPCR GPR52 (PDB: 6LI3), with the activating loop depicted in red.

**Fig. S9.**
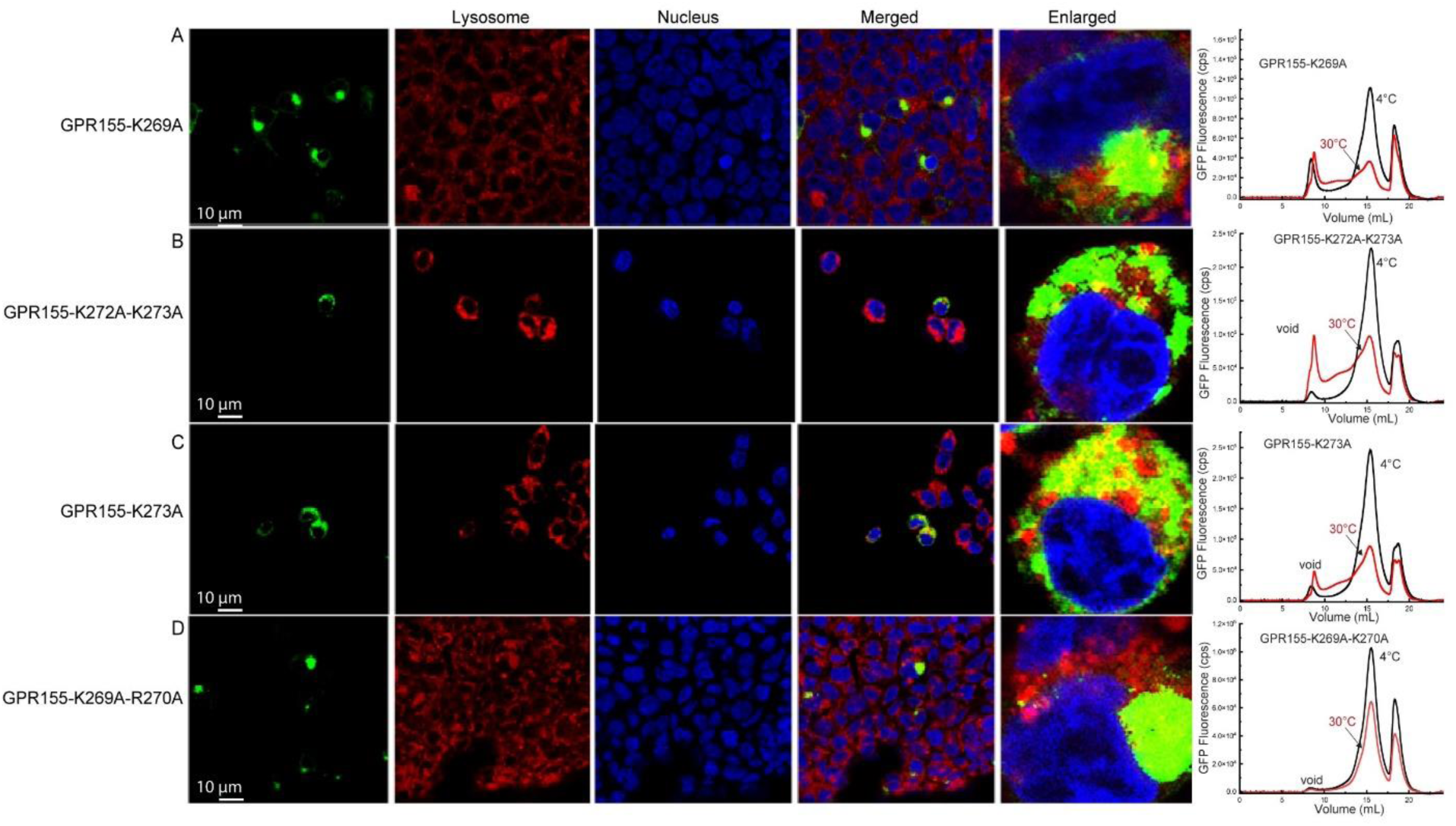
Mutational analysis of lipid binding residues. Confocal fuorescence imaging and thermostability of **(A)** GPR155-K269A, **(B)** GPR155-K272A-K273A, **(C)** GPR155-K273A, **(D)** GPR155-K269A-R270A

**Fig. S10.**
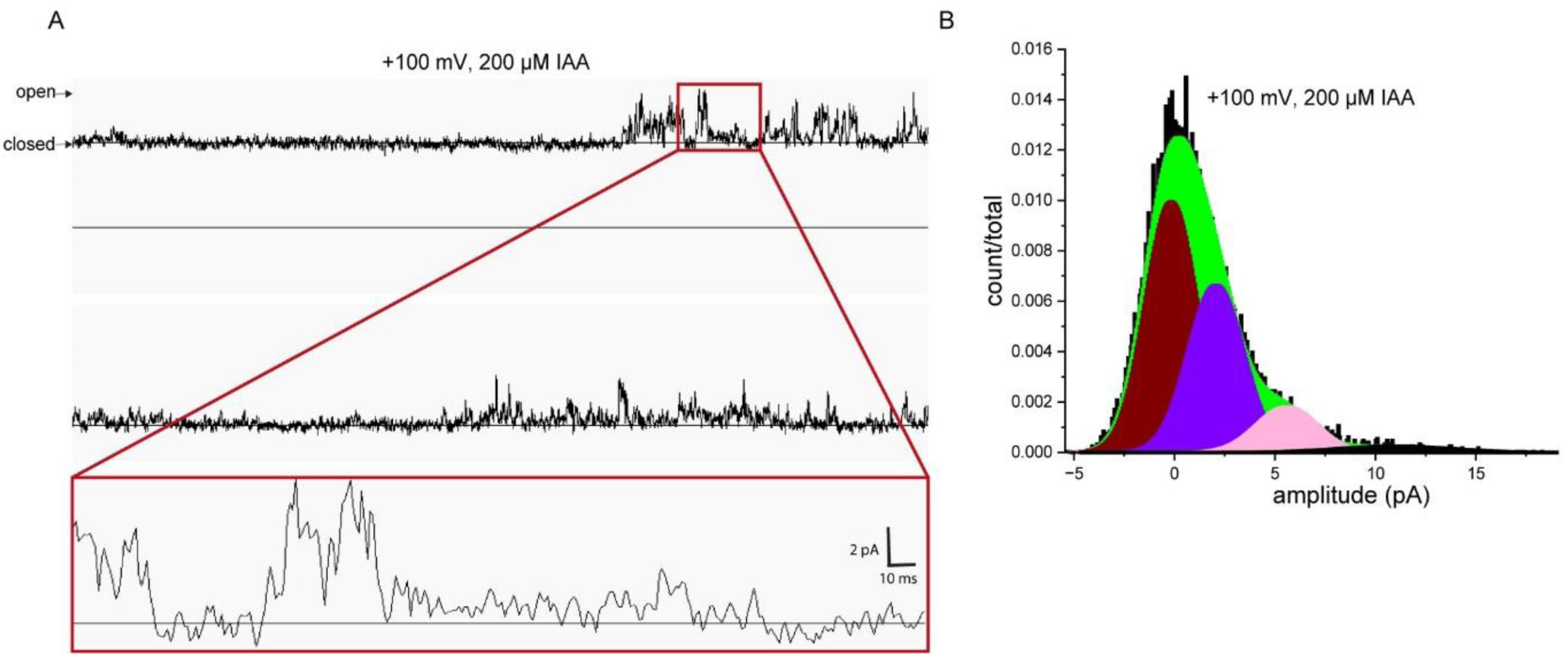
Lipid bilayer analysis of IAA transport by purified GPR155 in LMNG detergent.

**Table S1.** Cryo-EM data collection, refinement, and validation statistics.

|  | GPR155<sub>monomer</sub><br>(EMDB-xxxx)<br>(PDB xxxx) | GPR155<sub>PIN-like-dimer</sub><br>(EMDB-xxxx)<br>(PDB xxxx) | GPR155<sub>cholesterol-driven</sub><br><sub>dimer</sub> (EMDB-xxxx)<br>(PDB xxxx) |
| --- | --- | --- | --- |
| **Data collection and processing** |  |  |  |
| Voltage (kV) | 300 | 300 | 300 |
| Electron exposure (e<sup>-</sup>/Å<sup>2</sup>) | 48 | 48 | 48 |
| Defocus range (μm) | -1.6 to -2.5 | -1.6 to -2.5 | -1.6 to -2.5 |
| Pixel size (Å) | 1.1 | 1.1 | 1.1 |
| Symmetry imposed | C1 | C2 | C2 |
| Initial particle images (no.) | 6,757,838 | 6,757,838 | 6,757,838 |
| Final particle images (no.) | 151,034 | 135,237 | 123,102 |
| Map resolution (Å) | 3.4 | 3.8 | 3.95 |
| FSC threshold |  |  |  |
| Map resolution range (Å) | 2.6 to 7.0 | 2.6 to 7.0 | 2.6 to 7.0 |
| **Refinement** |  |  |  |
| Initial model used (PDB code) | This study | This study | This study |
| Model resolution (Å) | 3.4 | 3.8 | 3.95 |
| FSC threshold |  |  |  |
| Model resolution range (Å) | 2.6 to 7.0 | 2.6 to 7.0 | 2.6 to 7.0 |
| Map sharpening *B* factor (Å<sup>2</sup>) | -189 | -145 | -192 |
| Model composition |  |  |  |
| Non-hydrogen atoms | 4,688 | 9,537 | 9,378 |
| Protein residues | 585 | 1,162 | 1164 |
| Ligands | 6 | 14 | 10 |
| *B* factors (Å<sup>2</sup>) |  |  |  |
| Protein | 181.72 | 124.77 | 184.77 |
| Ligand | 145.54 | 132.33 | 165.33 |
| R.m.s. deviations |  |  |  |
| Bond lengths (Å) | 0.005 | 0.004 | 0.007 |
| Bond angles (°) | 1.05 | 0.94 | 1.08 |
| Validation |  |  |  |
| MolProbity score | 2.04 | 2.23 | 2.03 |
| Clashscore | 8.9 | 11.5 | 11.8 |
| Poor rotamers (%) | 0 | 0 | 0 |
| Ramachandran plot |  |  |  |
| Favored (%) | 94.13 | 91.51 | 95.32 |
| Allowed (%) | 5.53 | 7.99 | 4.42 |
| Disallowed (%) | 0.35 | 0.5 | 0.26 |

## References

1. S. T. Yang, A. J. B. Kreutzberger, J. Lee, V. Kiessling, L. K. Tamm, The role of cholesterol in membrane fusion. Chem Phys Lipids 199, 136–143 (2016).

2. X. Huang et al., Genome-Wide Analysis of the PIN Auxin Efflux Carrier Gene Family in Coffee. Plants (Basel) 9, (2020).

3. K. L. Ung et al., Structures and mechanism of the plant PIN-FORMED auxin transporter. Nature 609, 605-+ (2022).

4. Z. S. Yang et al., Structural insights into auxin recognition and efflux by PIN1. Nature 609, 611-+ (2022).

5. N. N. Su et al., Structures and mechanisms of the auxin transporter PIN3 (vol 609, pg 616, 2022). Nature 610, E2–E2 (2022).

6. J. D. Cohen, L. C. Strader, An auxin research odyssey: 1989-2023. Plant Cell 36, 1410–1428 (2024).

7. C. Xue et al., Tryptophan metabolism in health and disease. Cell Metab 35, 1304–1326 (2023).

8. Y. Zhou et al., The role of the indoles in microbiota-gut-brain axis and potential therapeutic targets: A focus on human neurological and neuropsychiatric diseases. Neuropharmacology 239, (2023).

9. D. W. Lamming, L. Bar-Peled, Lysosome: The metabolic signaling hub. Traffic 20, 27–38 (2019).

10. S. Inpanathan, R. J. Botelho, The Lysosome Signaling Platform: Adapting With the Times. Front Cell Dev Biol 7, (2019).

11. V. K. Mony, S. Benjamin, E. J. O’Rourke, A lysosome-centered view of nutrient homeostasis. Autophagy 12, 619–631 (2016).

12. H. R. Shin et al., Lysosomal GPCR-like protein LYCHOS signals cholesterol sufficiency to mTORC1. Science 377, 1290–1298 (2022).

13. S. Trifonov et al., GPR155: Gene organization, multiple mRNA splice variants and expression in mouse central nervous system. Biochem Biophys Res Commun 398, 19–25 (2010).

14. S. Trifonov et al., Lateral regions of the rodent striatum reveal elevated glutamate decarboxylase 1 mRNA expression in medium-sized projection neurons. Eur J Neurosci 35, 711–722 (2012).

15. C. Brochier et al., Quantitative gene expression profiling of mouse brain regions reveals differential transcripts conserved in human and affected in disease models. Physiol Genomics 33, 170–179 (2008).

16. X. C. Wang, Z. Liu, L. H. Jin, Anchor negatively regulates BMP signalling to control Drosophila wing development. Eur J Cell Biol 97, 308–317 (2018).

17. S. Umeda et al., Downregulation of GPR155 as a prognostic factor after curative resection of hepatocellular carcinoma. BMC Cancer 17, 610 (2017).

18. D. Shimizu et al., GPR155 Serves as a Predictive Biomarker for Hematogenous Metastasis in Patients with Gastric Cancer. Sci Rep 7, 42089 (2017).

19. H. J. Schulten et al., Comparison of microarray expression profiles between follicular variant of papillary thyroid carcinomas and follicular adenomas of the thyroid. Bmc Genomics 16, (2015).

20. E. Hacker et al., Reduced expression of IL-18 is a marker of ultraviolet radiation-induced melanomas. Int J Cancer 123, 227–231 (2008).

21. J. W. Lee et al., Genetic Characteristics Associated With Drug Resistance in Lung Cancer and Colorectal Cancer Using Whole Exome Sequencing of Cell-Free DNA. Front Oncol 12, 843561 (2022).

22. G. Diallinas, Transceptors as a functional link of transporters and receptors. Microb Cell 4, 69–73 (2017).

23. G. Van Zeebroeck, B. M. Bonini, M. Versele, J. M. Thevelein, Transport and signaling via the amino acid binding site of the yeast Gap1 amino acid transceptor. Nature Chemical Biology 5, 45–52 (2009).

24. S. A. Burchett, Regulators of G protein signaling: A bestiary of modular protein binding domains. J Neurochem 75, 1335–1351 (2000).

25. D. R. Ballon et al., DEP-domain-mediated regulation of GPCR signaling responses. Cell 126, 1079–1093 (2006).

26. D. N. Patil et al., Cryo-EM structure of human GPR158 receptor coupled to the RGS7-Gbeta5 signaling complex. Science 375, 86–91 (2022).

27. E. Jeong, Y. Kim, J. Jeong, Y. Cho, Structure of the class C orphan GPCR GPR158 in complex with RGS7-Gbeta5. Nat Commun 12, 6805 (2021).

28. C. Hunte et al., Structure of a Na/H antiporter and insights into mechanism of action and regulation by pH. Nature 435, 1197–1202 (2005).

29. E. Padan, H. Michel, NhaA: A Unique Structural Fold of Secondary Active Transporters. Isr J Chem 55, 1233–1239 (2015).

30. C. Lee et al., A two-domain elevator mechanism for sodium/proton antiport. Nature 501, 573-+ (2013).

31. Y. L. Dong et al., Structure and mechanism of the human NHE1-CHP1 complex. Nat Commun 12, (2021).

32. A. M. Duran, J. Meiler, Inverted topologies in membrane proteins: a mini-review. Comput Struct Biotechnol J 8, e201308004 (2013).

33. P. Becker et al., Mechanism of substrate binding and transport in BASS transporters. Elife 12, (2023).

34. E. Padan, Functional and structural dynamics of NhaA, a prototype for Na and H antiporters, which are responsible for Na and H homeostasis in cells. Bba-Bioenergetics 1837, 1047–1062 (2014).

35. N. J. Hu, S. Iwata, A. D. Cameron, D. Drew, Crystal structure of a bacterial homologue of the bile acid sodium symporter ASBT. Nature 478, 408–411 (2011).

36. S. Vosolsobě, K. Kurtović, V. Schmidt, R. Skokan, J. Petrášek, Origin and evolution of Auxin Efflux Carrier family: PIN, PILS, GPR155 and the others. bioRxiv, 2024.2006.2026.600818 (2024).

37. S. R. M. Bennett, J. Alvarez, G. Bossinger, D. R. Smyth, Morphogenesis in Pinoid Mutants of Arabidopsis-Thaliana. Plant J 8, 505–520 (1995).

38. V. Cherezov et al., High-resolution crystal structure of an engineered human beta2-adrenergic G protein-coupled receptor. Science 318, 1258–1265 (2007).

39. K. Palczewski et al., Crystal structure of rhodopsin: A G protein-coupled receptor. Science 289, 739–745 (2000).

40. A. Bortolato et al., Structure of Class B GPCRs: new horizons for drug discovery. Br J Pharmacol 171, 3132–3145 (2014).

41. F. Y. Siu et al., Structure of the human glucagon class B G-protein-coupled receptor. Nature 499, 444–449 (2013).

42. X. Lin et al., Structural basis of ligand recognition and self-activation of orphan GPR52. Nature 579, 152–157 (2020).

43. N. Hoppe et al., GPR161 structure uncovers the redundant role of sterol-regulated ciliary cAMP signaling in the Hedgehog pathway. Nat Struct Mol Biol 31, 667–677 (2024).

44. X. Lin et al., Cryo-EM structures of orphan GPR21 signaling complexes. Nat Commun 14, 216 (2023).

45. Z. Yang et al., Structure of GPR101-Gs enables identification of ligands with rejuvenating potential. Nat Chem Biol 20, 484–492 (2024).

46. A. Goehring et al., Screening and large-scale expression of membrane proteins in mammalian cells for structural studies. Nat Protoc 9, 2574–2585 (2014).

47. K. Saotome, A. K. Singh, M. V. Yelshanskaya, A. I. Sobolevsky, Crystal structure of the epithelial calcium channel TRPV6. Nature 534, 506–511 (2016).

48. C. J. Russo, L. A. Passmore, Electron microscopy: Ultrastable gold substrates for electron cryomicroscopy. Science 346, 1377–1380 (2014).

49. S. H. Scheres, RELION: implementation of a Bayesian approach to cryo-EM structure determination. J Struct Biol 180, 519–530 (2012).

50. A. Punjani, J. L. Rubinstein, D. J. Fleet, M. A. Brubaker, cryoSPARC: algorithms for rapid unsupervised cryo-EM structure determination. Nature methods 14, 290 (2017).

51. T. Bepler et al., Positive-unlabeled convolutional neural networks for particle picking in cryo-electron micrographs. Nat Methods 16, 1153–1160 (2019).

52. E. F. Pettersen et al., UCSF Chimera--a visualization system for exploratory research and analysis. Journal of computational chemistry 25, 1605–1612 (2004).

53. P. Emsley, B. Lohkamp, W. G. Scott, K. Cowtan, Features and development of Coot. Acta Crystallogr D Biol Crystallogr 66, 486–501 (2010).

54. P. V. Afonine et al., Towards automated crystallographic structure refinement with phenix.refine. Acta Crystallogr D 68, 352–367 (2012).

55. T. D. Goddard et al., UCSF ChimeraX: Meeting modern challenges in visualization and analysis. Protein Sci 27, 14–25 (2018).

56. E. C. Meng et al., UCSF ChimeraX: Tools for structure building and analysis. Protein Sci 32, e4792 (2023).

57. P. V. Afonine et al., Towards automated crystallographic structure refinement with phenix.refine. Acta Crystallogr D Biol Crystallogr 68, 352–367 (2012).

58. O. Klykov, S. P. Gangwar, M. V. Yelshanskaya, L. Yen, A. I. Sobolevsky, Structure and desensitization of AMPA receptor complexes with type II TARP gamma5 and GSG1L. Mol Cell 81, 4771–4783 e4777 (2021).

59. L. S. Milescu, C. L. Nicolai, F. Qin, F. Sachs, New developments in the QUB software for single-channel data analysis. Biophysical Journal 82, 267a–267a (2002).

60. L. Milescu, A. Galick, F. Qin, A. Auerbach, F. Sachs, New software in the QUB suite. Biophysical Journal 76, A208–A208 (1999).

61. F. Qin, Restoration of single-channel currents using the segmental k-means method based on hidden Markov modeling. Biophys J 86, 1488–1501 (2004).

62. T. J. Collins, ImageJ for microscopy. Biotechniques 43, 25–30 (2007).

